# A broadly active fucosyltransferase *LmjFUT1* whose mitochondrial localization and catalytic activity is essential in the parasitic protozoan *Leishmania*

**DOI:** 10.1101/2021.05.10.443387

**Authors:** Hongjie Guo, Sebastian Damerow, Luciana Penha, Stefanie Menzies, Gloria Polanco, Hicham Zegzouti, Michael A. J. Ferguson, Stephen M. Beverley

## Abstract

Glycoconjugates play major roles in the infectious cycle of the trypanosomatid parasite *Leishmania*. While GDP-Fucose synthesis is essential (Guo *et al* 2017), fucosylated glycoconjugates have not been reported in *Leishmania major*. Four predicted fucosyltransferases appear conventionally targeted to the secretory pathway; *SCA1/2* play a role in side-chain modifications of lipophosphoglycan, while gene deletion studies here showed that *FUT2* and *SCAL* were not essential. Unlike most eukaryotic glycosyltransferases, the predicted α 1-2 fucosyltransferase encoded by *FUT1* localized to the mitochondrion. A quantitative ‘plasmid segregation’ assay, expressing *FUT1* from the multicopy episomal pXNG vector in a chromosomal null Δ*fut1*^-^ background, established that *FUT1* is essential. Similarly “plasmid shuffling” confirmed that both enzymatic activity and mitochondrial localization were required for viability, comparing import-blocked or catalytically inactive enzymes respectively. Enzymatic assays of tagged proteins expressed *in vivo* or of purified recombinant FUT1 showed it had a broad fucosyltransferase activity including glycan and peptide substrates. Unexpectedly a single rare Δ*fut1*^-*s*^ segregant (*Δfut1^s^*) was obtained in rich media, which showed severe growth defects accompanied by mitochondrial dysfunction and loss, all of which were restored upon *FUT1* re-expression. Thus, *FUT1* along with the similar *Trypanosoma brucei* enzyme TbFUT1 (Bandini *et al* 2021) joins the eukaryotic O-GlcNAc transferase isoform as one of the few glycosyltransferases acting within the mitochondrion. Trypanosomatid mitochondrial FUT1s may offer a facile system for probing mitochondrial glycosylation in a simple setting and their essentiality renders it an attractive target for chemotherapy of these serious human pathogens.

**Significance Statement:** Abundant surface glycoconjugates play key roles in the infectious cycle of protozoan parasites including *Leishmania*. Through defining biosynthetic pathways we identified a fucosyltransferase FUT1 that was localized to the parasite mitochondrion, an atypical compartment for glycosyltransferases. FUT1 was essential for normal growth, requiring both mitochondrial localization and enzymatic activity. Loss of FUT1 in a unique segregant showed extensive mitochondrial defects. Enzymatic tests showed FUT1 could fucosylate glycan and peptide substrates *in vitro*, although as yet the native substrate is unknown. Trypanosomatid mitochondrial FUT1s may offer a facile system in the future for probing mitochondrial glycosylation in a setting uncomplicated by multiple isoforms targeted to diverse compartments, and its essentiality renders it an attractive target for chemotherapy of these deadly parasites.

## Introduction

*Leishmania* is a widespread human pathogen, with more than 1.7 billion people at risk, several hundred million infected, and with 12 million showing active disease ranging from mild cutaneous lesions to severe disfiguring or ultimate lethal outcomes (2–4). The *Leishmania* infectious cycle alternates between extracellular promastigote in the midgut of sand flies and intracellular amastigote residing within macrophages of the mammalian host, where it survives and proliferates in highly hostile environments. These parasites have evolved specific mechanisms enabling them to endure these adverse conditions, including a dense cell surface glycocalyx composed of lipophosphoglycan (LPG), glycosylphosphatidylinositol (GPI) anchored proteins (GP63 and GP46), glycosylinositolphospholipids (GIPLs) and secreted glycoconjugates such as proteophosphoglycan (PPG) and secreted acid phosphatase (sAP) (reviewed in 5; 6-8). One prominent feature of LPG, PPGs and sAPs is the presence of disaccharide phosphate repeating units ([6Gal(β)1,4)Man(α1)-PO_4_]), also termed phosphoglycan or PG repeats.

Our lab has focused on both forward and reverse genetic approaches to map out glycoconjugate synthesis in *Leishmania*, emphasizing genes impacting the glycocalyx (9, 10) as well as ether lipids and sphingolipids (11, 12). These studies have provided powerful tools leading to new insights on the requirements for LPG and related phosphoglycan -bearing molecules in both parasite stages within the mammalian and sand fly hosts (6, 7, 13–19).

In several *Leishmania* species modifications of the dominant PG repeats of LPG and PPG play key roles in the insect stages in mediating both the attachment and release of promastigotes and metacyclics, respectively, from the sand fly midgut via binding to midgut receptors there (20). In *L. major* strain FV1 (LmjF), β1-3 galactosyl modifications of the PG repeating units enable replicating promastigotes to bind to the midgut lectin PpGalec, while addition of D-Arabinopyranose (D-Ara*p*) to the side-chain galactosyl residues block this interaction and allow release of parasites for subsequent transmission (20–22). Accordingly, we used forward genetic analysis to identify a large family of LPG side chain galactosyltransferases (SCG1-7) and D-Arabino*pyranosyl*transferases (SCA1/2) mediating these key modifications (22–27).

As D-Ara*p* is relatively uncommon in nature (28), we were motivated to explore its synthetic pathway. Previously we identified two genes showing strong homology to the bifunctional *Bacteroides* protein FKP mediating synthesis of GDP-L-Fucose (GDP-Fuc) through successive kinase and pyrophosphorylase steps (29, 30). Many enzymes using L-Fucose will also accept D-Ara*p* (which differ only by the 6-methyl group), and assays of the two recombinant *Leishmania* proteins showed that indeed one could synthesize both GDP-D-Ara*p* and GDP-Fuc (AFKP; LmjF.16.0480), while the second could only synthesize GDP-Fuc (FKP; LmjF.16.0440; (30)). These data were consistent with studies showing the presence of both GDP-Fuc and GDP-Ara*p* in *Leishmania* (31). Correspondingly, genetic studies showed that knockouts of *AFKP* completely abrogated GDP-Ara*p* and arabinosylated LPG synthesis, while knockouts of *FKP* showed little effect (30). However, we were unable to knockout both genes simultaneously, suggesting an unanticipated role for GDP-Fuc. That the essential role of A/FKPs depended on GDP-Fucose was established when GDP-Fuc but not GDP-Ara*p* was provided through expression of the two *de novo* GDP-Fucose enzymes from *Trypanosoma brucei*, GDP-mannose 4,6-dehydratase (GMD) and GDP-Fuc synthetase, also known as GDP-4-dehydro-6-deoxy-D-mannose epimerase/reductase (GMER) (32). Similarly the loss of de novo GDP-fucose synthesis was also lethal in *T. brucei*, which lacks the FKP salvage pathway (32).

The essentiality of GDP-Fuc was unexpected since there are few reports of fucosylated molecules in *Leishmania.* One is a “Fucose Mannose Ligand” from *L. donovani* (33) for which a definitive structure is lacking. Several proteins were predicted to be fucosylated from mass spectrometric proteome studies of *L. donovani*, albeit without experimental confirmation (34). More recently, human erythropoietin expressed in *L. tarentolae* was shown to bear fucosylated glycans (35). While untested, it seemed unlikely that any of these putative fucosylations would be essential for growth in culture. Together, these data suggest that any putative essential Fuc-glycoconjugate predicted from genetic deletion studies must be relatively rare and/or cryptic.

To better understand the fucose requirement in *Leishmania*, and guide efforts to identify the essential fucose-conjugate, we surveyed candidate fucosyltransferases (FUTs). Informatic studies revealed five candidate FUTs (Table 1), of which only *FUT1* was conserved in all trypanosomatids, and the only one essential in *Leishmania*. Unexpectedly, FUT1 localizes within the parasite mitochondrion, an uncommon observation for glycosyltransferases. Thus we obtained genetic evidence that this localization, as well as its catalytic activity, is essential for viability, and that its absence leads to severe mitochondrial dysfunction.

**Table 1.**
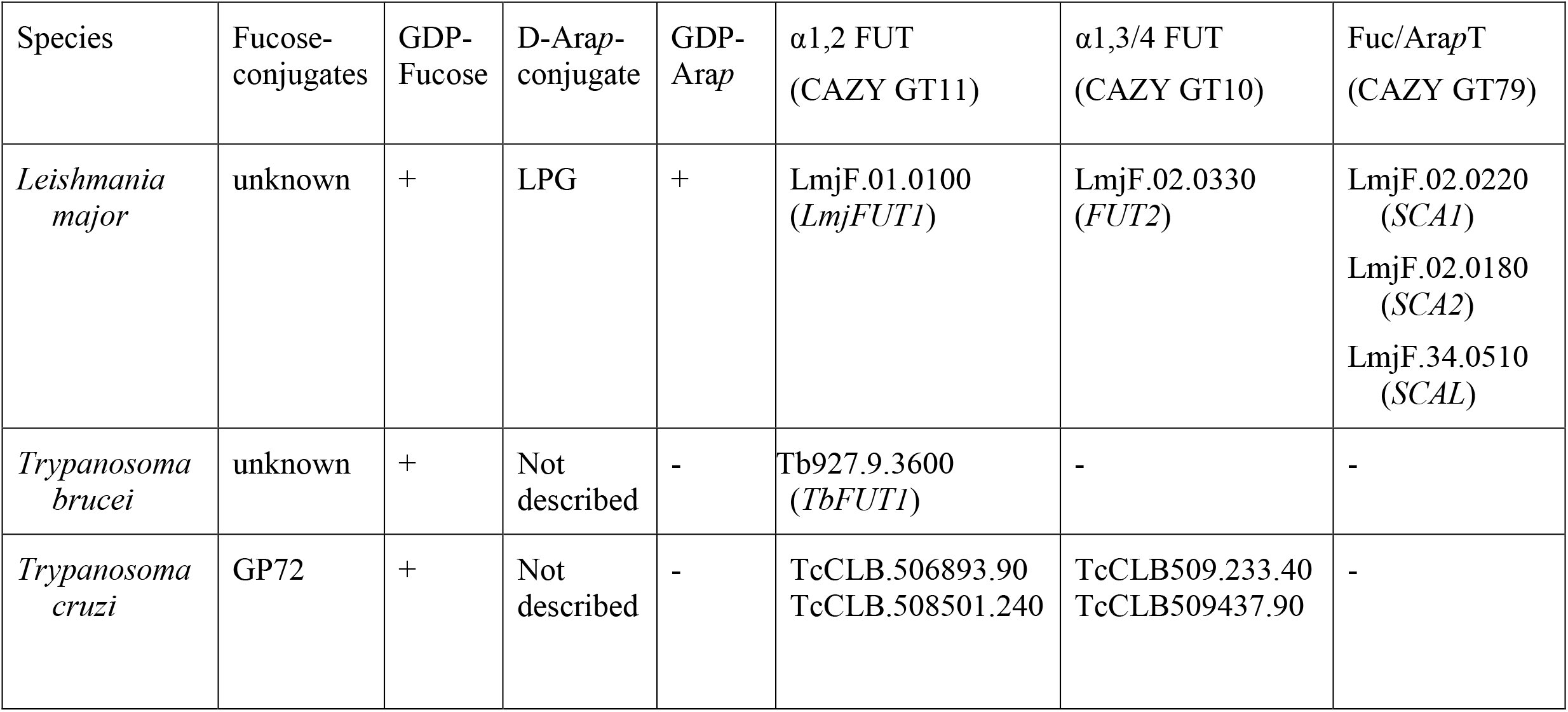
Trypanosomatid L-Fucose/D-Arabinosyl*pyranose* conjugates, precursors, and candidate transferases

## Results

### Database mining for fucosyltransferase (FUTs)/ arabinopyranosyltransferase(AraTs) candidates in *L. major*

Using a diverse collection of FUTs compiled from GenBank or CAZY databases (36), we searched the predicted *L. major* proteome for proteins showing significant similarities and/or conserved motifs. Two new candidates (*FUT1,* LmjF01.0100; *FUT2*, LmjF.02.0330) as well as three previously described genes emerged *(SCA1,* LmjF02.0220*; SCA2,* LmjF02.0180*; SCAL,* LmjF34.0510; Table 1). Of these, only *FUT1* was conserved in all trypanosomes, while *FUT2* occurred in *Trypanosoma cruzi* (Table 1).

#### SCA1/2

The closely related *SCA1/2* (Side-Chain-Arabinosyltransferase) genes (>99% identity) had been identified functionally and enzymatically as Golgi α 1,2 D-Ara*p* transferases modifying Gal-substituted PG repeating units of LPG in *L. major* during metacyclogenesis (24, 27). Interestingly, when provided with high levels of external L-Fucose the LPG PG repeating units can become fucosylated, likely mediated by these two enzymes (37). Since *Leishmania* require neither galactosylated nor arabinosylated LPG for growth in culture or metacyclogenesis (15, 38), their study was deprioritized here.

#### SCAL

(SCA1/2-like) was identified with 31% identity to the SCA1/2 proteins, which together constitute CAZY family 79 (27, 36). The lack of any related genes in *T. brucei* and *T. cruzi* is consistent with the absence of GDP-D-Ara*p* in both (31) (Table 1). We were able to knock-out both alleles of *SCAL* without difficulty and the mutant showed no appreciable growth defect in the culture (Fig S1), suggesting that this gene is not essential *in vitro*, or is redundant with other proteins.

#### FUT2

The predicted FUT2 protein was identified by similarity to the CAZY family GT10, which comprises α1,3 or 1,4 fucosyltransferases (36). Strong homology of the 1604 aa FUT2 protein was seen with PFAM family 00852 (α1,3 or 1,4 fucosyltransferases) from residues 1049-1159, along with other motifs characteristic of this family (39, 40) . Using a plasmid segregation test (described further below for *FUT1*), we were able to knockout both alleles of *FUT2* without difficulty (Fig S2), suggesting that this gene is not essential during *in vitro* culture, or is redundant with other proteins.

#### FUT1

The predicted FUT1 protein showed relationships to the CAZY GT11 family comprised mostly of α 1, 2 fucosyltransferases (36). Unlike the four other FUTs, orthologous genes were found in all three trypanosomatid genomes (Table 1). The annotated Lmj*FUT1* predicted a protein of 348 amino acids, exhibiting four characteristic motifs of the GT11 family (Fig. 1), of which motif I (HVRRGDY) has been implicated in the binding of GDP-fucose (40). While SCA1, SCA2, SCAL and FUT2 resembled typical eukaryotic fucosyltransferases, predicted to be type II membrane proteins within the secretory pathway (40, 41), no transmembrane domain was detected in FUT1. Instead, FUT1 surprisingly bore a predicted mitochondrial targeting sequence by two algorithms (88%; TargetP and MitoProt) (42, 43). For *L. major*, the predicted N-terminal mitochondrial targeting peptide (MTP) comprised the N-terminal 23 amino acids upstream of motif IV, rich in Arg and hydrophobic amino acids (Fig. 1). The LmjFUT1 mitochondrial localization and its requirement for viability were established experimentally below.

**Figure 1.**
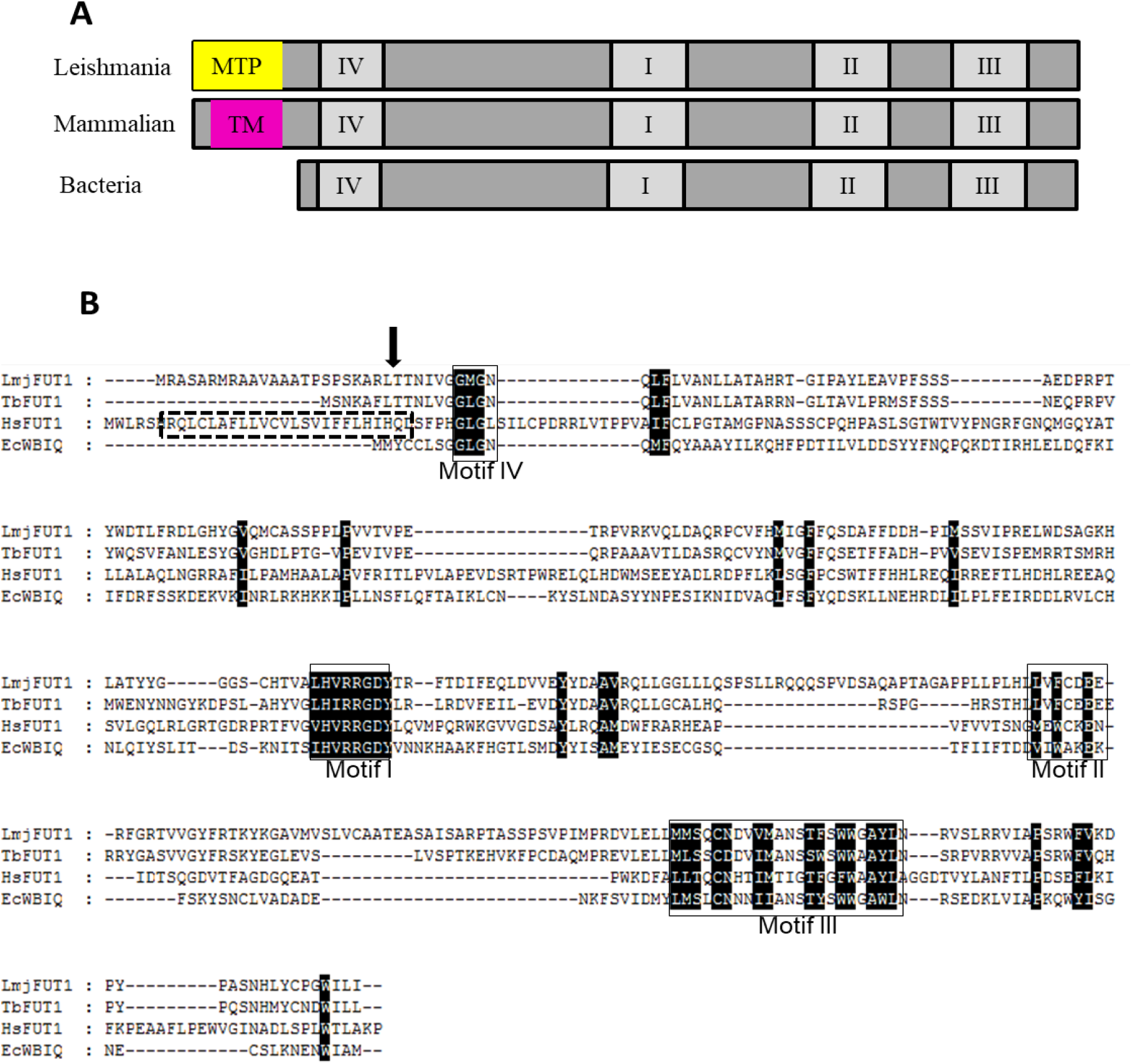
Comparison of mitochondrial *Leishmania major FUT1* relative to related fucosyltransferases. Panel A. Schematic of *Leishmania* FUT1 relative to mammalian or bacterial α1.2 fucosyltransferases, depicting conserved motifs I-IV, the mitiochondrial targeting peptide of LmjFUT1 (MTP, yellow) and the transmembrane domain of human FUT1 (TM, purple) relative to *E. coli* WBIQ. Panel B. Multiple sequence alignment of FUT1 and fucosyltransferases from *Trypanosoma brucei* (TbFUT1 from *T. brucei*), bacteria (EcWBIQ from E. coli O127) and mammals (human hsFUT1).. Identical or highly conserved residues are highlighted in black. Conserved fucosyltransferase motifs are marked by black boxes (40). The predicted mitochondrial peptide cleavage site is marked by an arrow. The predicated transmembrane domain for HsFUT1 is marked by a dash dot box.

### Generation of chromosomal null mutants (*Δfut1^-^*) in the presence of ectopically expressed *FUT1*

Anticipating that *FUT1* could encode an essential fucosyltransferase, we employed a plasmid segregation-based strategy, in which a counter-selectable ectopic copy of *FUT1* was introduced prior to ablation of the chromosomal *FUT1* alleles (44). WT *L. major* parasites were first transfected with an episomal pXNGPHLEO-*FUT1* construct, followed by successive inactivation of the two chromosomal *FUT1* alleles using *SAT* and *PAC* targeting constructs. PCR tests confirmed that the marker replacements occurred as planned and the chromosomal alleles of *FUT1* were precisely deleted, with retention of the episomal FUT1 (Fig. 2A,B; Fig S3). For simplicity chromosomal deletions are hereafter referred to as Δ*fut1*^-^, with this line termed Δ*fut1*^-^ / +pXNGPHLEO-*FUT1*.

**Figure 2.**
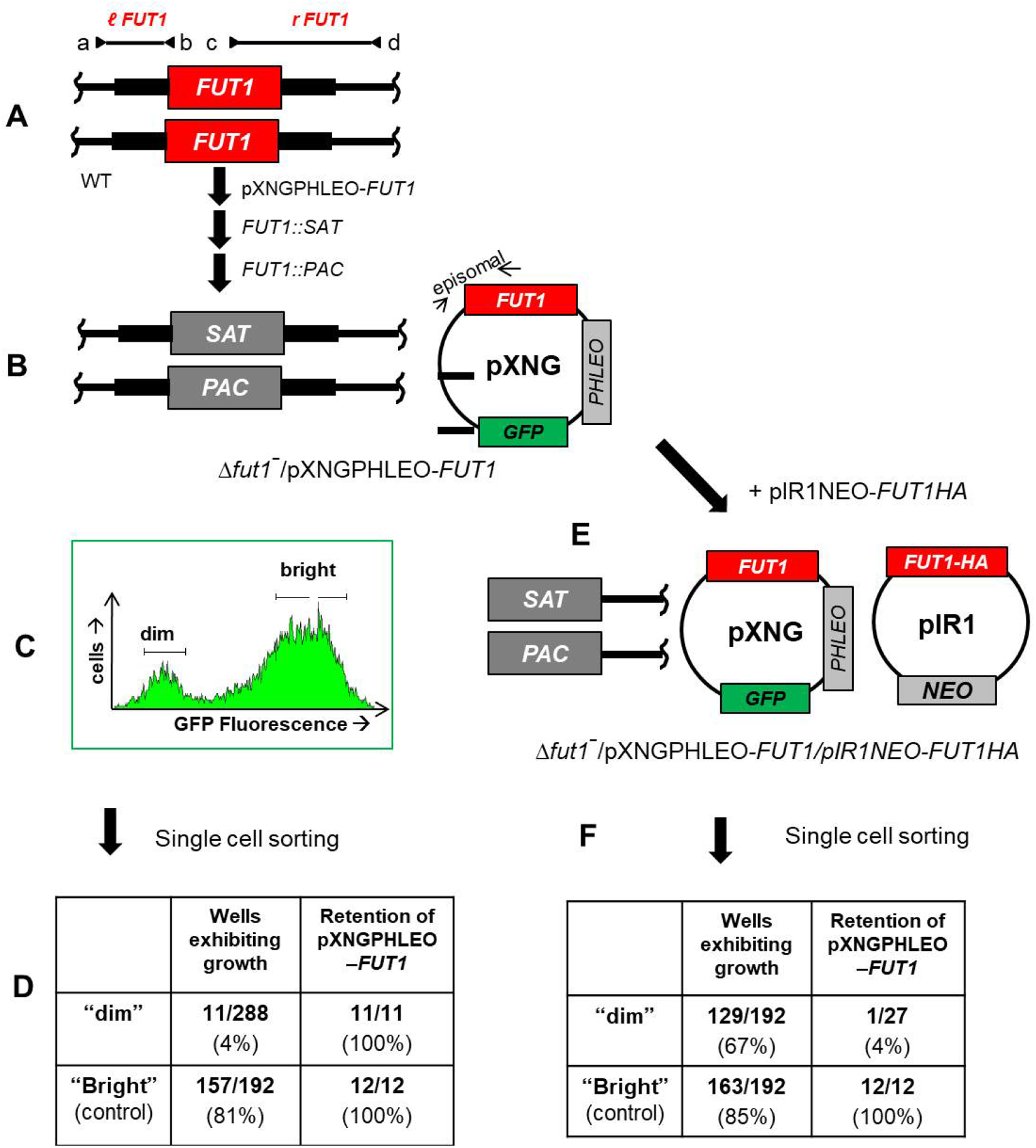
Plasmid segregation tests show *FUT1* is essential. Panel A. *FUT1* locus and transfection. The figure shows the WT FUT1 locus, with the ORF as a central red box and the flanking sequences used for homologous replacement by SAT or PAC drug resistance cassettes (FUT1::SAT or FUT1::PAC respectively) shown as heavier flanking lines. The location of ‘outside’ primers used to establish planned replacements (a/b or c/d; to confirm replacements SAT or PAC specific primers were substituted for b or c; Fig. S3) are shown. WT *L. major* was first transfected with pXNGPHLEO-FUT1, and successively with the FUT1::SAT and FUT::PAC constructs. Panel B. The chromosomal null Δ*fut1***^-^***^-^* mutant obtained in the presence of episomal *FUT1*. The genoetype of this line is FUT::ΔSAT/FUT1::ΔPAC/ +pXNGPHLEO-FUT1, abbreviated as Δ*fut1***^-^***/+*pXNGPHLEO-FUT1. PCR confirmation tests of the predicted genotype are found in Fig. S3. pXNG additionally expresses GFP (44). Panel C. Plasmid segregation tests of FUT1 essentiality. Δ*fut1***^-^***/ +*pXNGPHLEO-FUT1 was grown for 24 h in the absence of Phleomycin, and analyzed by GFP flow cytometry. The gates used for subsequent quantitation and/or sorting, weakly-fluorescent (dim) and fluorescent (bright) parasites are as shown. Panel D. Results of sorting experiment shown in panel C. Single cells from both GFP dim or bright populations sorted into 96 well plates containing M199 medium, and incubated for 2 weeks, at which time growth was assessed. The numbers sorted and their growth and properties are shown, with ‘bright’ cells mostly surviving and ‘dim’ cells mostly not . All 11 survivors from the ‘dim’ and 12 tested from the ‘bright’ population sort were confirmed to retain of pXNGPHLEO-FUT1 by growth in media containing phleomycin. Panel E. Generation of line required for plasmid shuffling. Δ*fut1***^-^***/ +*pXNGPHLEO-FUT1 was transfected with pIR1NEO-FUT1-HA expressing C-terminally tagged FUT1. PCR tests confirming the predicted genotype Δ*fut1***^-^***/ +*pXNGPHLEO-FUT1/ +pIR1NEO-FUT1-HA are shown in Fig. S3). This line was then grown 24 hr in the absence of phleomycin and subjected to cell sorting and growth tests as described in panel C. Panel F. Results of sorting experiment described in panel E. In this experiment, both ‘dim’ and ‘bright’ cells mostly survived (67% and 85% respectively). 27 of the ‘dim’ cells were tested of which 26 had lost pXNGPHLE-FUT1 by PCR tests, thereby representing the desired Δ*fut1***^-^**/ +pIR1NEO-FUT1-HA.

For plasmid segregation tests, parasites were grown briefly (24 h) in the absence of phleomycin to allow loss of pXNGPHLEO. Parasites were then analyzed for GFP expression by flow cytometry, as a measure of pXNG copy number (44). Two populations emerged: a large population of ‘bright’ cells showing strong fluorescence bearing high copy numbers of pXNGPHLEO-*FUT1*(>200 FU); and a smaller population of ‘dim’ cells, exhibiting control/background fluorescence levels (2-20 FU), lacking pXNG-*FUT1* completely, or bearing only a few copies (Fig. 2C).

Fluorescence-activated cell sorting was then used to recover single cells into individual wells of a 96 well microtiter plate, containing M199 medium without phleomycin. For the ‘bright’ cell population, 157/192 cells inoculated with single “bright” parasites grew out (81%), representing the ‘cloning/plating’ efficiency of cells subjected to this protocol. In contrast, growth was seen in only 11/288 of the ‘dim’ parasites tested similarly (4 %; Fig. 2D). The 11 survivors still retained pXNGPHLEO-*FUT1* as judged by their ability to grow out in the presence of phleomycin and expression of GFP (data not shown), likely arising by imperfect sorting or other technical factors as seen previously (44). After correction for the plating/cloning efficiency, we estimated that *FUT1*-null mutants were not obtained from 236 cells tested. This greatly exceeds the resolution of classic ‘failed to recover deletion’ protocols and suggests that under these conditions, *FUT1* is essential.

### FUT1 is localized in mitochondria

To confirm the surprising prediction of mitochondrial localization, we expressed a C-terminal HA-tagged FUT1 protein using the pIR1NEO vector as an episome; this construct was introduced into the Δ*fut1^-^ /* +pXNGPHLEO*-FUT1* line, yielding Δ*fut1^--^/* +pXNGPHLEO*-FUT1* / pIR1NEO-*FUT1*-HA (Fig. 2E). We then used “plasmid shuffling” to select for a line which had lost untagged pXNG-*FUT1* construct but maintained the tagged pIR-*FUT1.* Following single cell sorting, 85% of the bright cell wells grew out (Fig. 2F). Similarly, 67 % of the dim cells also grew out, and subsequent tests showed that 96% (26/27) tested lacked GFP and were phleomycin sensitive, while retaining tagged FUT1-HA, yielding Δ*fut1^-^/*+ pIR1NEO-*FUT1*-HA (Fig. 2F). These data confirmed that a C terminal HA tag did not comprise *FUT1* function.

Immunofluorescence analysis (IFA) was used first to assess localization of HA-tagged FUT1 in the Δ*fut1^-^ /* +pIR1NEO-*FUT1*-HA line. Anti-HA antibody was used to stain the parasites after labeling with mitochondrial marker MitoTracker Red, and the co-localization coefficient was calculated (Pearson’s). As anticipated from the initial predication, the HA-tagged FUT1 co-localized completely with the mitochondrial marker with a Pearson’s co-localization index of 0.97 ± 0.03 (Fig. 3A).

**Figure 3.**
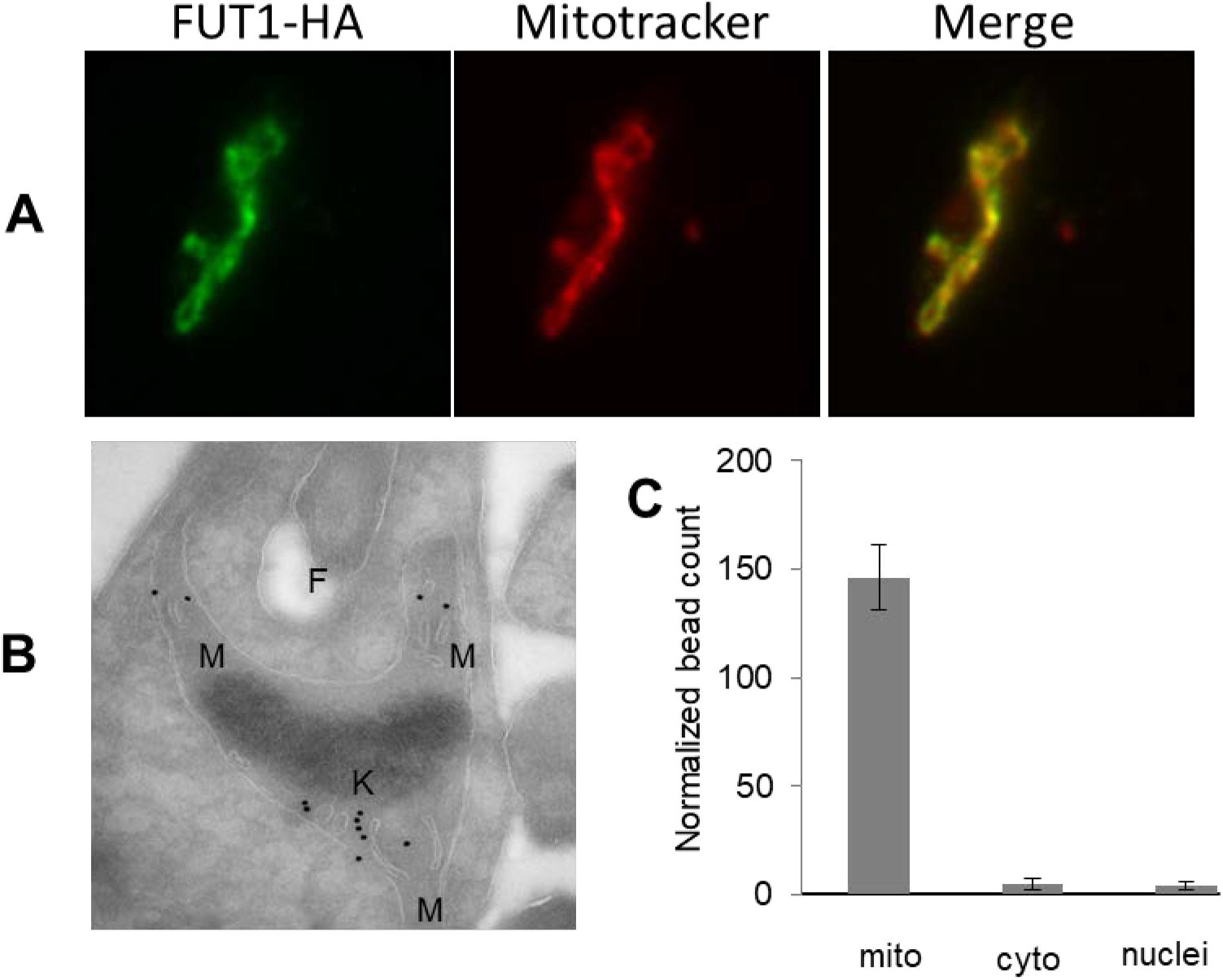
LmjFUT1 is localized to the parasite mitochondrion. Panel A. Indirect immunofluorescence of parasites lines of Δ*fut1***^-^**/+ pIR1NEO-FUT1-HA incubated with anti-HA antibody (leftmost image, green), Mitotracker Red CMXRos (central image, red), and merged (rightmost image, yellow) are shown. Colocalization by Pearson’s r coefficientwas 0.97. Panel B. Cryo-immuno-EM with anti-HA beads of Δ*fut1***^-^**/+ pIR1NEO-FUT1-HA. M, mitochondrion; K, kinetoplast DNA network; F, flagellar pocket. Panel C. Quantitation of cryo-immuno-EM anti-HA bead labelling to FUT-HA in cellular compartments. The data shown are the bead counts and standard deviation, taken from 3 different experiments comprising 3 sets of 10 sections each, and normalized to the relative compartment area.

Cryo-immunoelectron microscopy using anti-HA conjugated beads binding to FUT-HA showed binding primarily to the mitochondrion, generally within the lumen (Fig. 3B). Quantitation of bead density showed more than 94% binding to the mitochondrion, far greater than seen to other compartments (Fig. 3C). Similar results were obtained with C-terminal HA-tagged FUT1 expressed in WT cells. In contrast, expression of an N-terminally HA tagged FUT1 in WT parasites failed to give any signal in western blotting or IFA using anti-HA antibody, consistent with the prediction that N-terminal FUT1 functions as a mitochondrial targeting peptide, which would be expected to be cleaved off during import. We tested deletions of the extended LmjFUT1 MTP region for mitochondrial localization and/or function (Fig. S4). A 16 aa truncation (MSKAR) was fully functional and mitochondrially targeted, a 17 aa truncation (MKAR) was partially functional and mitochondrially targeted, and a 18 aa truncation (MAR) was not functional but partially mitochondrially targeted (Fig S4). These data suggest first that LmjFUT1 may possess a cryptic mitochondrial targeting sequence, resembling the N terminus of TbFUT1, which we show elsewhere is also mitochondrially targeted (1). These data further suggest that in trypanosomatids critical residues of the Motif IV region may extend functionally further towards the N terminus than in other species.

### Recombinant FUT1 is an active fucosyltransferase

When expressed in *E. coli* most FUT1 was insoluble, despite trials with several protein fusion expression systems, a common observation for eukaryotic glycosyltransferases (45) (Fig. 4A, lanes 1-4). The best results were often GST-FUT1 fusions, yielding a small fraction of soluble recombinant protein (<5%), which could be purified by GST affinity chromatography (Fig. 4A, lanes 5,6). The fusion protein had an apparent molecular weight of 66 kDa similar to the theoretical value (64 960 Da), and the major impurity appeared to be GST (Fig 4A, lane 6). All preparations tended to be unstable, and assays were performed on freshly purified material including an active acceptor control (below). Attempts to cleave the GST tag from the fusion protein using Factor Xa yielded little free FUT1, and thus the fusion protein was used for enzymatic characterizations.

**Figure 4.**
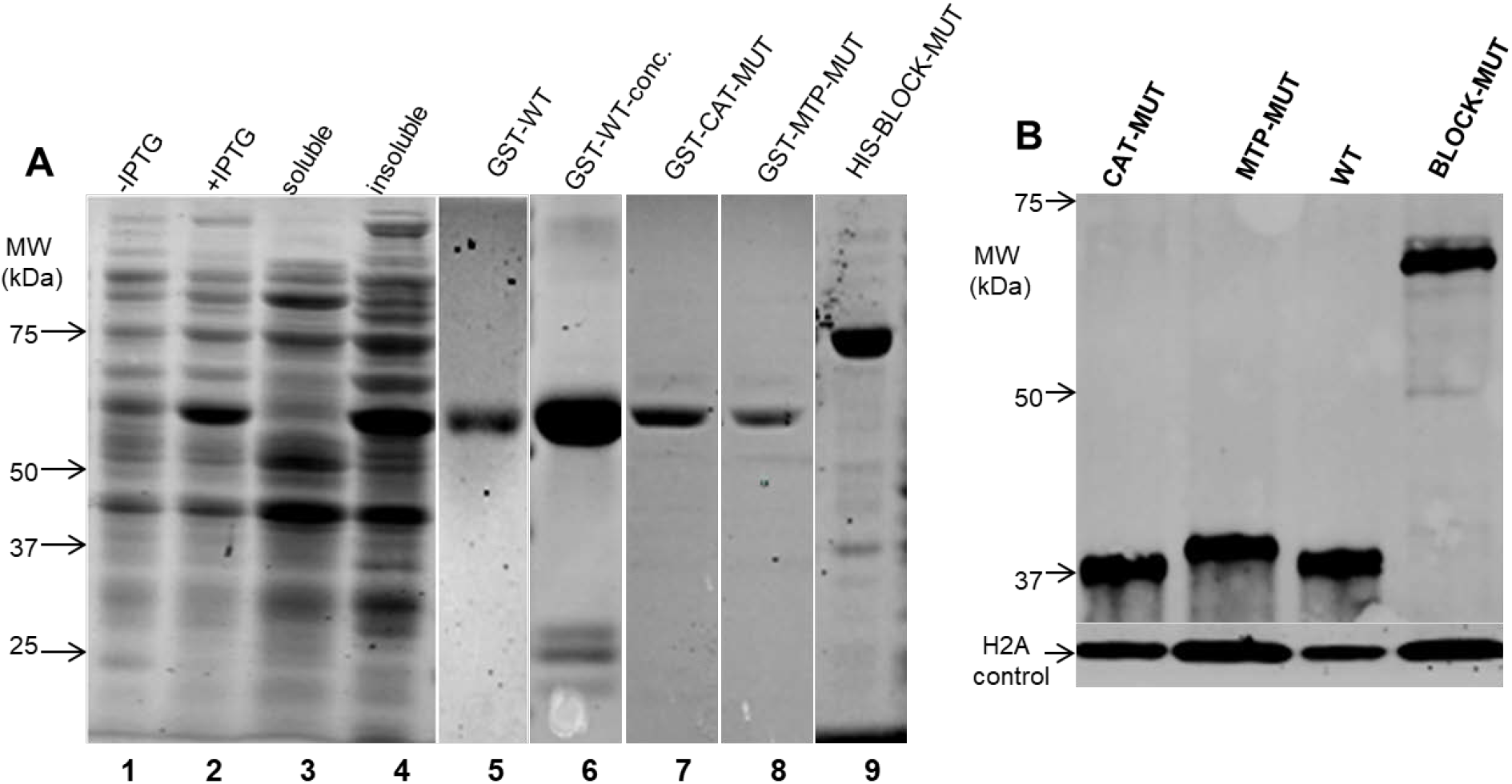
Expression of WT or mutant FUT1s in *E. coli* or within *Leishmania.* Panel A: Expression and purification of recombinant GST-FUT1 fusion proteins from *E. coli* visualized follows SDS-PAGE. Lanes 1-6 are GST-FUT1 (WT). Lane 1: before-induction whole cell lysates; lane 2: post-induction whole cell lysates; lane 3: soluble fraction; lane 4: insoluble fraction; lane 5: elution; lane 6: concentrated eluted protein. Lane 7 Purified GST-CAT-MUT; Lane 8, purified GST-MTP-MUT; and lane 9, purified HIS-BLOCK-MUT. Molecular weight Markers are shown on the left. The images from lanes 1-4 and 5,6,7, 8 and 9 are from separate experiments. Panel B. Western blot analysis of C-terminal HA-tagged FUT1s expressed in *Leishmania*. Lysates from parasites expressing the indicated HA-tagged FUT1s in a Δ*fut1****^-^*** /+ pXNGPHLEO-FUT1 background are shown (the presence of untagged WT FUT1 was required as none of the mutants were viable in its absence; Fig 6). Western blots were performed with anti-HA to visualized the tagged FUT1 expression, and anti-*L. major* H2A as a loading control

To assay recombinant GST-FUT1 we used an indirect assay which measures GDP arising from glycosyltransferase activity, through coupling to ATP synthesis and measurement by a luciferase/luciferin reaction (46). Initially, recombinant GST-FUT1 was incubated with GDP-Fuc as donor and β-D-Gal-(1→3)-D-GlcNAc (lac-N-biose; LNB) as acceptor. A reaction time course established the product to increase linearly for the first 6 min and to plateau by 20 min (Fig 5A).

**Figure 5.**
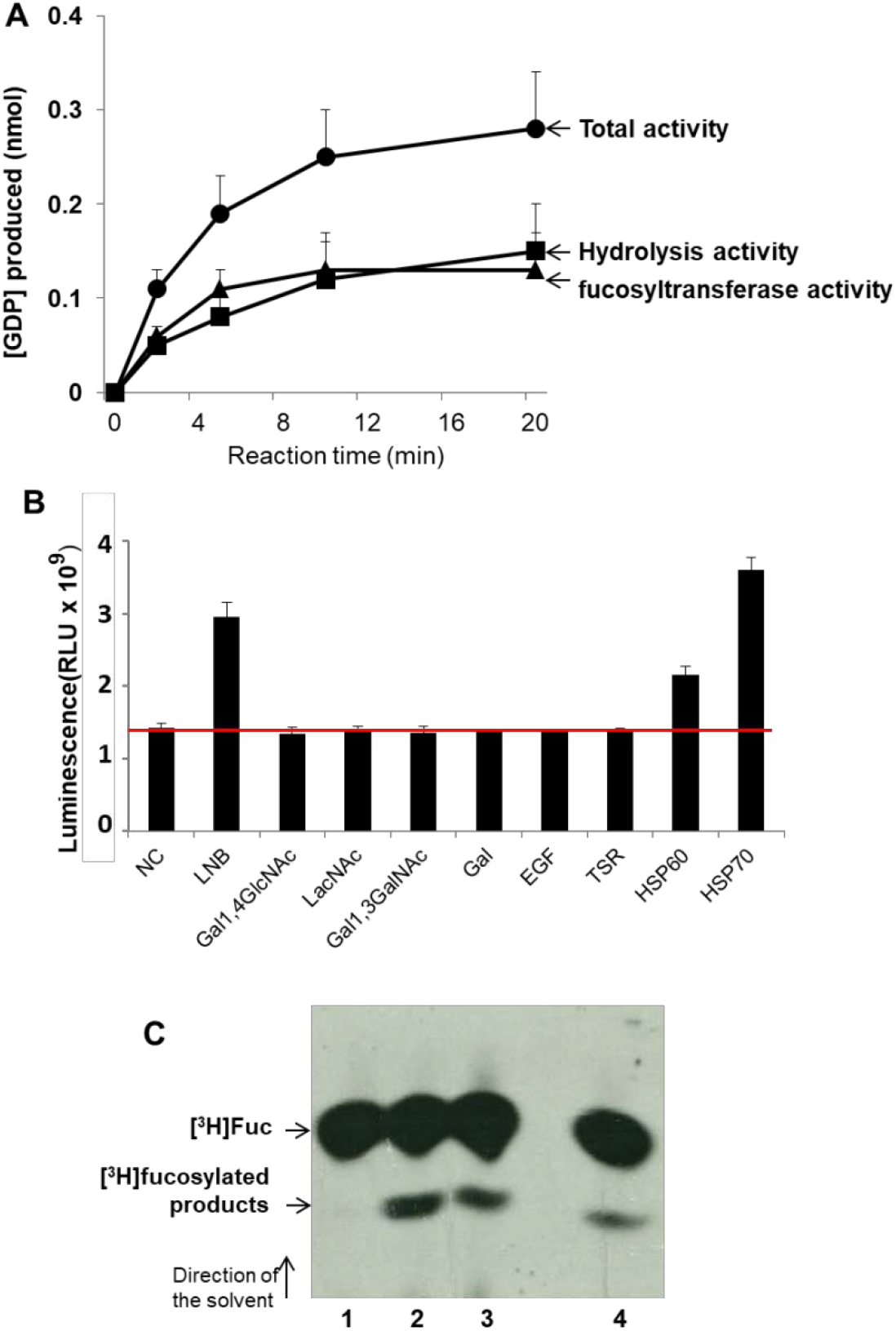
LmjFUT1 is a fucosyltransferase. Panel A: Time course of GDP production with GST-FUT1 in the presence or absence of GDP-Fucose and 1 mM LNB. Solid circles, incubation in the presence of GDP-Fucose + LNB (“total activity”); solid squares, incubation with GDP-Fucose alone (hydrolysis activity); solid triangles, difference between presence and absence of LNB (acceptor dependent fucosyltransferase activity). Panel B. GST-FUT1 was incubated with GDP-Fucose in the presence of various acceptors, and the luminescence from GDP produced was measured after 5 min of incubation. Lane 1 NC, no acceptor (hydrolysis only); lane 2, LNB; lane 3, Gal1,4GlcNAc; lane 4, LacNAc; Lane 5, Gal1,3GalNAc; lane 6; Galactose; lane 7, EGF, EGF-like repeat; lane 8, thrombospondin like repeat; lane 9, mHSP60 peptide; and lane 10, mHSP70 peptide. The red horizontal line represents the hydrolysis activity (lane 1). Panel C: Fucosyltransferase activity assayed in *L. major in vivo.* Whole cell lysates were incubated with anti-HA agarose beads, which were washed, and then incubated with 1μCi GDP-[^3^H]-Fuc and 1 mM LNB as acceptor. Reaction products were separated by HPTLC and detected by fluorography as shown. Lane 1, WT; lane 2, Δ*fut1***^-^**/ + pIR1NEO-LmjFUT1-HA; lane 3, Δ*fut1***^-^***/ +*pIR1NEO-TbFUT1; lane 4, WT / + pIR1NEO-Lmj-FUT1-HA.

In the absence of the acceptor, GDP-Fuc was hydrolyzed and an increase in GDP could be detected (Fig 5A, curve labelled ‘hydrolysis activity’). Such acceptor independent activity has been described previously for other FUTs in the absence or with sub-optimal substrates (46, 47). In the presence of LNB acceptor GDP formation increased ∼2-fold (Fig. 5B, curve labelled “Total activity”). Acceptor-dependent fucosyltransferase activity was calculated from the difference between these two curves (Fig 5A; curve labelled “fucosyltransferase activity”). From these findings initial rate data were calculated from a 5 min reaction, and established that the optimum conditions for the coupled reaction were 37°C, pH 6 with LNB as an acceptor, yielding a specific activity of recombinant GST-FUT1 of 59 ± 5.4 nmol/min/mg protein (Fig. 6E). In these assays, recombinant LmjFUT1 lacked activity with the glycan acceptors N-acetyllactosamine (LacNAc), β-Galactose, Galβ1,3GalNAc (Fig. 5B). Overall the activity seen with the substrates tested mirrored those observed elsewhere for TbFUT1, where a wider range was tested but with LNB similarly being most active (1).

**Figure 6.**
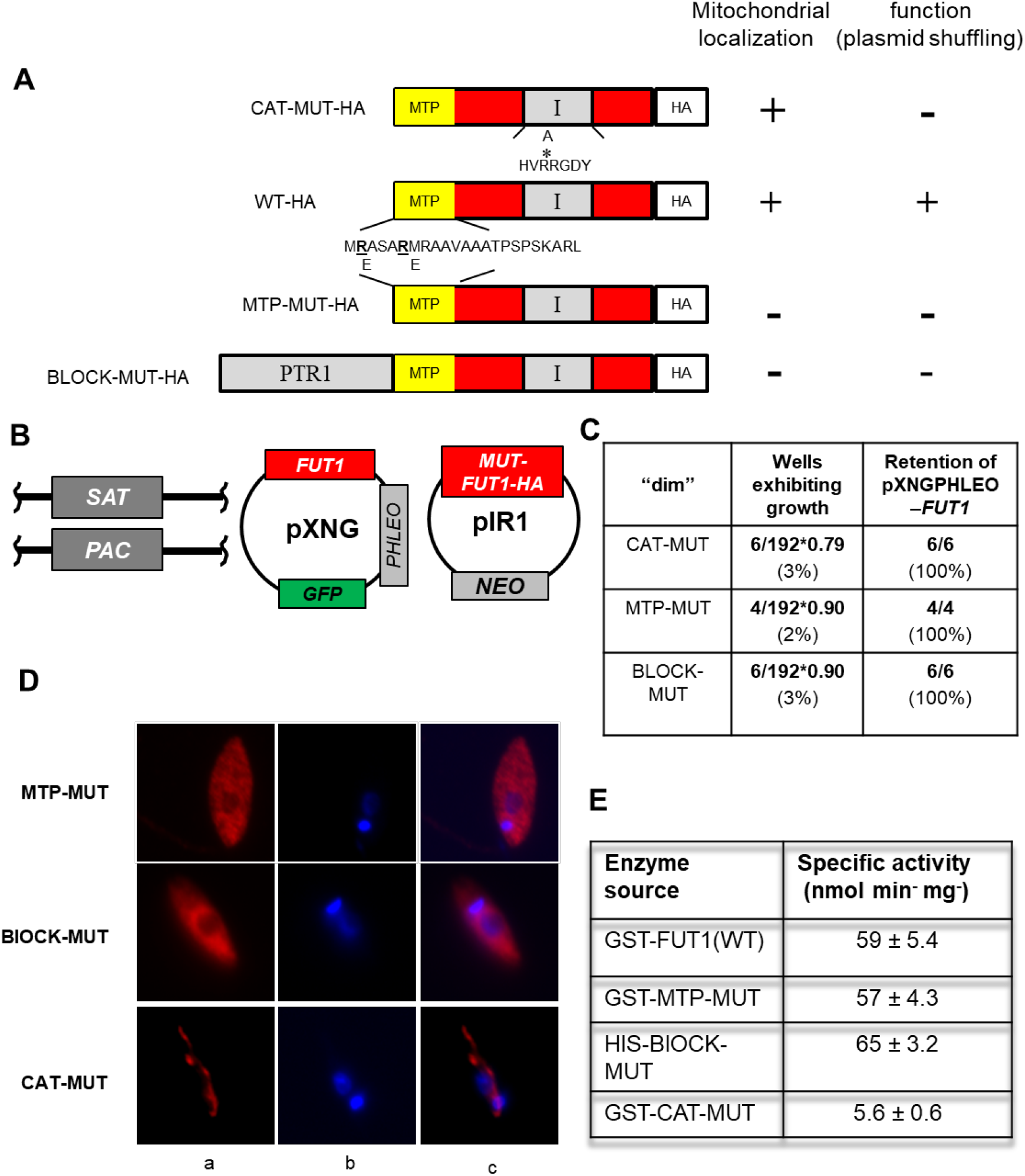
Both the localization in mitochondrion and fucosyltransferase activity are required for the essential function of FUT1. Panel A. Depiction of mutant LmjFUT1-HA designed to block mitochondrial import or catalysis. The mitochondrial targeting peptide (MTP) is shown in yellow, the catalytic motif I is shown in gray, and the HA tag is shown in white, with the remainder of FUT1 in red. CAT-MUT-HA has an R297A substitution; MTP-MUT-HA replaces two Glu residues in the MTP with Arg; BLOCK-MUT-HA has the cytoplasmic protein PTR1 fused to the N-terminus. The results of mitochondrial localization tests (Panel D) and plasmid shuffling tests (Panel C) are summarized on the right of the illustrations. Panel B. Scheme of plasmid shuffling to test the function of mutant FUT1s. FUT1s were expressed from pIR1NEO (“pIR1NEO-MUT-FUT-HA”) in the Δ*fut1***^-^***/ +*pXNGPHLEO-FUT1 line. Growth in the absence of phleomycin and FACS sorting of ‘dim’ and ‘bright’ cells was performed as described in Fig. 2. Panel C. Plasmid shuffling tests of FUT1 mutants. Δ*fut1***^-^**pXNG-FUT1/pIR-MUT-HA lines was grown for 24 h in the absence of phleomycin, and analyzed by GFP flow cytometry. In all tests, survival of “bright” (control” cells was 79-90%. The results show that few dim cells yielded growth, and all of those arose from incomplete sorting (retention of pXNGPHLEO-FUT1) Panel D: Indirect immunofluorescence of HA tagged FUT1 expressed from pIRNEO in the Δ*fut1***^-^***/+*pXNGPHLEO-FUT1 background. Column a, anti-HA (red); column b, DNA (Hoechst, blue), and column c, merge of columns a and b. Panel E. Acceptor-dependent specific activity of purified recombinant FUT1 proteins assayed by GDP formation in the presence of GDP-Fucose and LNB. The average and standard deviation of three preparations is shown.

We confirmed the fucosyltransferase activity of recombinant FUT using enzyme expressed within *Leishmania*. Whole cell lysates of WT or *Δfut1**^-^*** parasites bearing pIR1NEO-FUT1-HA were applied to anti-HA-agarose beads, which were washed and then incubated with GDP-[^3^H]-Fuc and LNB. With both cell lines, a new product was observed, similar to that seen with a control expressing TbFUT1, shown elsewhere to be to Fucα1,2Galβ1,3GlcNAc (1). As expected, no products other than [^3^H]-Fuc were found with WT lacking tagged FUT1, again attributed to hydrolysis of the GDP-Fuc substrate (Fig. 5A,C). As the *Δfut1**^-^*** pIR1NEO-TbFUT1 mutant was obtained by plasmid shuffling in a *Δfut1**^-^*** background (Experimental Procedures), these data proved this heterologous enzyme could satisfy the *Leishmania* FUT1 requirement (Fig 5C, lane 3).

Two peptide substrates for mammalian protein-O-transferases 1 and 2, corresponding to the epidermal growth factor-like repeats (EGFR) or thrombospondin-like repeats (TSR; kindly provided by R. Haltiwanger; (48, 49)) lacked activity with recombinant LmjFUT1 (Fig. 5B). We synthesized peptides bearing 5 amino acids on either side of the proposed fucosylation sites of the *L. infantum* mitochondrial HSP60 or HSP70 (34), which both showed significant acceptor-dependent fucosyltransferase activity (Fig. 5B). To confirm this unexpected activity against a peptide substrate, we incubated LmjFUT1 with the mHSP70 peptide and GDP-Fucose, and analyzed the fucosylated product by mass spectrometry, where fucosylation of the central target serine residue was clearly evident (Fig. S5). Thus, LmjFUT1 could recognize a diverse spectrum of acceptors *in vitro*, including both glycan and peptide substrates.

### Mitochondria localization is required for the essential role(s) of FUT1

While FUT1 was found primarily in the mitochondrion (Fig. 3), occasionally the predominant distribution of a protein may be misleading about its true site of function (50). Thus, we asked whether mitochondrial localization was required for *FUT1* function, by testing mutants where the mitochondrial targeting was compromised (Fig. 6). Based on characteristic features of mitochondrial targeting peptides (MTPs; (51)), we replaced the first two Arg residues with Glu, predicted to reduce the probability of mitochondrial import from 88% to 3% (MitoProtII; MTP-MUT, Fig 6A). Similarly, we fused the cytosolic protein PTR1 to the N-terminus of FUT1 (BLOCK-MUT), anticipating that the large 30 kDa PTR1 would block recognition of a far-internal MTP (Fig. 6A). Both modifications were made with the C-terminally HA tagged FUT1 expressed in pIR1NEO, rendering them suitable for immune-detection and functional tests (Fig. 6B). Plasmid shuffling tests showed that neither MTP-MUT nor BLOCK-MUT could substitute for WT-FUT1, as only 2-3% of the ‘dim’ cells survived (vs. 90 ∼ 90 % for ‘bright’ cells; Fig 6C). Again, tests showed that the few surviving dim cells had retained the WT FUT1 plasmid and thus were not true segregants (Fig. 6C).

Immunofluorescence analysis of the tagged proteins showed that unlike the HA-tagged WT FUT1, HA-tagged MTP-MUT and BLOCK-MUT proteins were now localized to the cytoplasm (Fig 6D). Two essential controls were performed to validate the conclusion above. First, western blot analysis showed that despite their cytosolic localization, expression of the MTP-MUT or BLOCK-MUT proteins was comparable to that of the WT FUT1 in *Leishmania*, relative to an H2A loading control (Fig, 4B). Consistent with the cytosolic localization and lack of mitochondrial targeting peptide cleavage, the size of MTP-MUT-HA was slightly greater than WT-HA, calculated to be about 40 kDa, close to the theoretical uncut protein (39.5 kDa; Fig 4B). Second, we expressed and purified tagged MTP-MUT and BLOCK-MUT proteins in *E. coli*, and showed that their specific FUT activity was the same as WT (Fig 6E). Thus, the failure of the mutants to rescue the chromosomal Δ*fut1**^-^*** mutant did not arise because of the failure to express an active *FUT1*, but instead arose solely from their failure to enter the mitochondrion.

### Fucosyltransferase activity is required for the essential role(s) of FUT1

We next tested whether the essential function of FUT1 required fucosyltransferase activity, perhaps through a structural role (52). We mutated the first arginine to alanine (R297A) in the FUT1 motif I (Fig. 1B), conserved in many fucosyltransferase families (40) and mutation of which causes 97-100% loss of in catalytic activity (53, 54). A C-terminal HA tagged catalytic FUT1 mutant (CAT) was expressed using the pIR1NEO vector, and immunofluorescence assays showed it remained in the mitochondrion like WT (Fig 6D). However, when subjected to plasmid shuffling only 3 % of the ‘dim’ cells grew out, vs 79% for ‘bright’ cells (Fig 6C). Subsequent tests of the ‘dim’ survivors showed they still retained the WT *FUT1* plasmid (Fig. 6C).

These data suggested that CAT nutant was unable to support *Leishmania* growth. Again, two controls strongly support this interpretation. First, western blot analysis showed that the level of CAT mutant expression within *Leishmania* was comparable to that of a tagged WT FUT1, relative to an H2A loading control (Fig. 4B). As expected, the size of CAT-HA was similar to WT FUT1-HA, indicating that it was similarly cleaved during import (Fig. 4B). Secondly, we expressed and purified the CAT mutant protein from *E. coli*, and assayed FUT activity. While the behavior and yield during purification of this enzyme were similar to the WT and MTP mutants, the CAT-MUT only retained less than 10% of WT activity, very close to the background of this assay (Fig. 6E).

Collectively, these data argue that expression of an enzymatically active mitochondrion fucosyltransferase is required for *Leishmania* viability.

### Recovery of a rare *Δfut1^s^* mutant by plasmid segregation in rich medium

The segregation tests above yielding no *FUT1*-null cells were performed in standard *Leishmania* M199 media (Fig. 2). Reasoning that a richer media might spare mitochondrial deficiencies arising from *FUT1* loss, we repeated this experiment several times using a richer medium, Schneider insect medium. In multiple experiments collectively screening more than 1000 cells, few “dim” cells survived as before (Fig 7B), even after extended incubations. However, in just one experiment a ‘dim’ clone was recovered which completely lacked *FUT1* by PCR tests (Fig. 7A-C). When both WT and the sole *fut1-*null mutant (here termed *Δfut1**^s^**)* were inoculated into fresh media, WT cells doubled after about 8 h, while *Δfut1**^s^*** increased slowly in cell number for the first 2 weeks, after which it grew somewhat more rapidly and eventually reached similar cell density as WT (Fig. 7D). This pattern of very slow initial growth repeated in subsequent passages, although after extended passage the growth increased somewhat. The growth defect of primary *Δfut1**^s^*** could be fully restored by restoration of FUT1 expression (Fig. 7D), as could all other defects described below.

**Figure 7.**
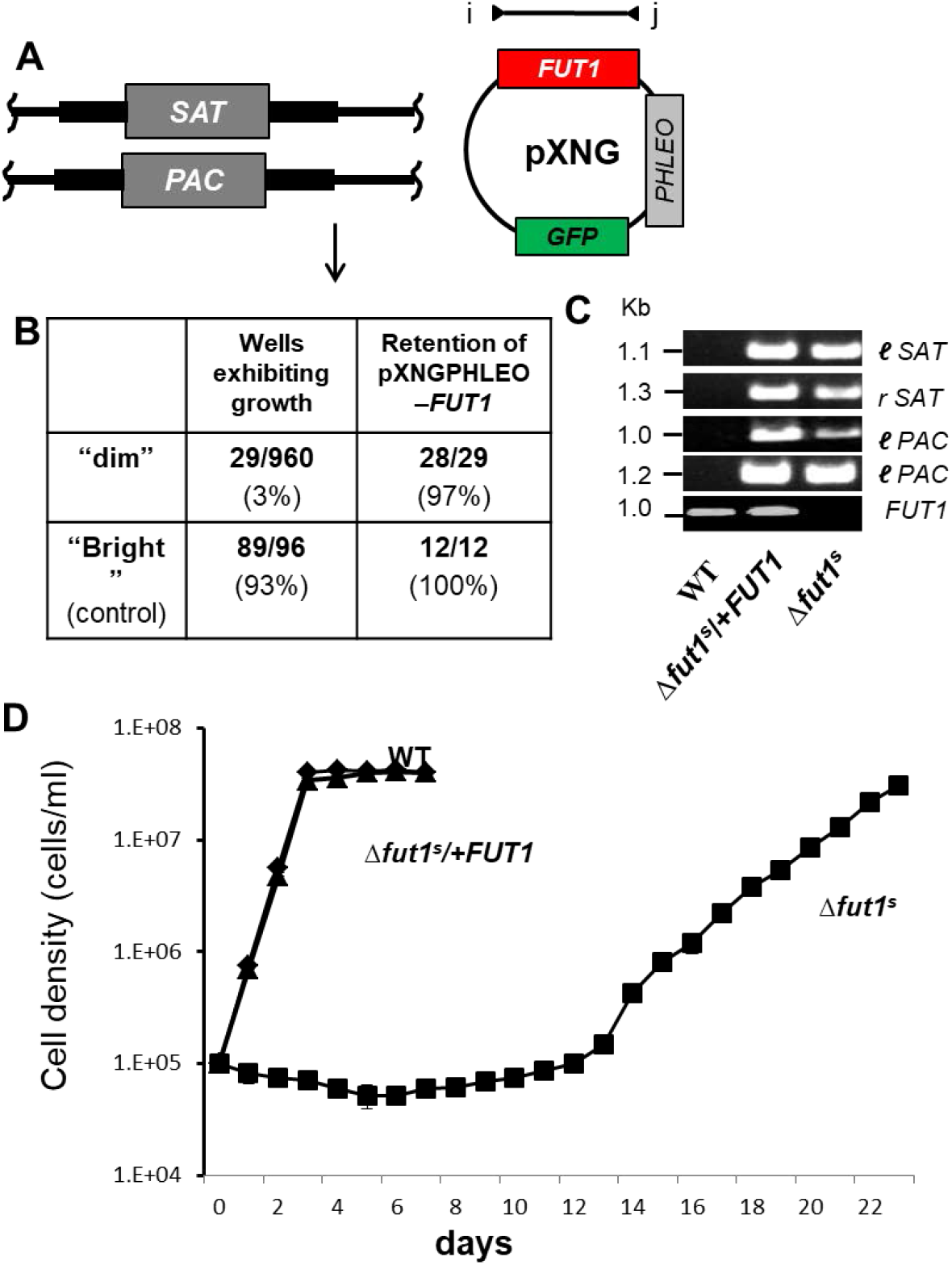
Recovery of a single *FUT1*-null mutant segregant (Δ*fut1^s^*). Panel A. Starting line for plasmid segregation in rich culture medium. Δ*fut1***^-^***/ +*pXNGPHLEO-FUT1 cells were grown and subjected to single cell sorting in Schneider’s media. Panel B. Control “bright” cells showed a typically high frequency of wells supporting growth (93%), while few “dim” cells survived as in M199 medium (Fig. 2). Of the 29 survivors, 28 retained the pXNGPHLEO plasmid, while one designated Δ*fut1****^s^*** did not. Panel C. PCR tests of Δ*fut1****^s^***, WT and a Δ*fut1****^s^*** /pXNGPHLEO-FUT1 ‘addback”. Primer sets l-SAT, r-SAT, l-PAC and r-PAC confirm Δ*fut1****^s^*** lacks chromosomal FUT1 (Fig. S3) and FUT1 ORF primers (l, 3955; j, 3956) confirm the absence of *FUT1* sequences in Δ*fut1****^s^*** and its presence in the addback. Panel D. Growth of WT, Δ*fut1****^s^*** and Δ*fut1****^s^***/+pXNGPHLEO-FUT1 in M199 medium. Parasites were inoculated at a density of 10^5^/ml and growth was followed by coulter counting.

### *Δfut1^s^* showed ultrastructural and functional mitochondrial changes

Transmission EM of *Δfut1**^s^*** revealed changes in the ultrastructure of mitochondria, with swelling and bloated cristae (Fig. 8A) often containing dark electron dense aggregates, similar to that seen previously in disrupted cristae associated with dysfunctional mitochondria (55). The kDNA network showed alterations in size, in *Δfut1**^s^*** decreasing 80% in length while increasing 12% in thickness relative to WT (Figs. 8B, S6). DAPI staining to reveal nuclear and kinetoplast DNA revealed that while WT parasites mostly showed a typical 1 kinetoplast – 1 nuclei (1K1N) pattern (56) only 42% of *Δfut1**^s^*** were 1K1N, with 39% 1K2N; remarkably, 18% lost kDNA entirely (0K1N; Fig 8C, D). Few abnormalities were observed in the ultrastructure of other organelles such as the nucleus, endoplasmic reticulum, flagellum, and acidocalcisomes (Fig. 8A,B).

**Figure 8.**
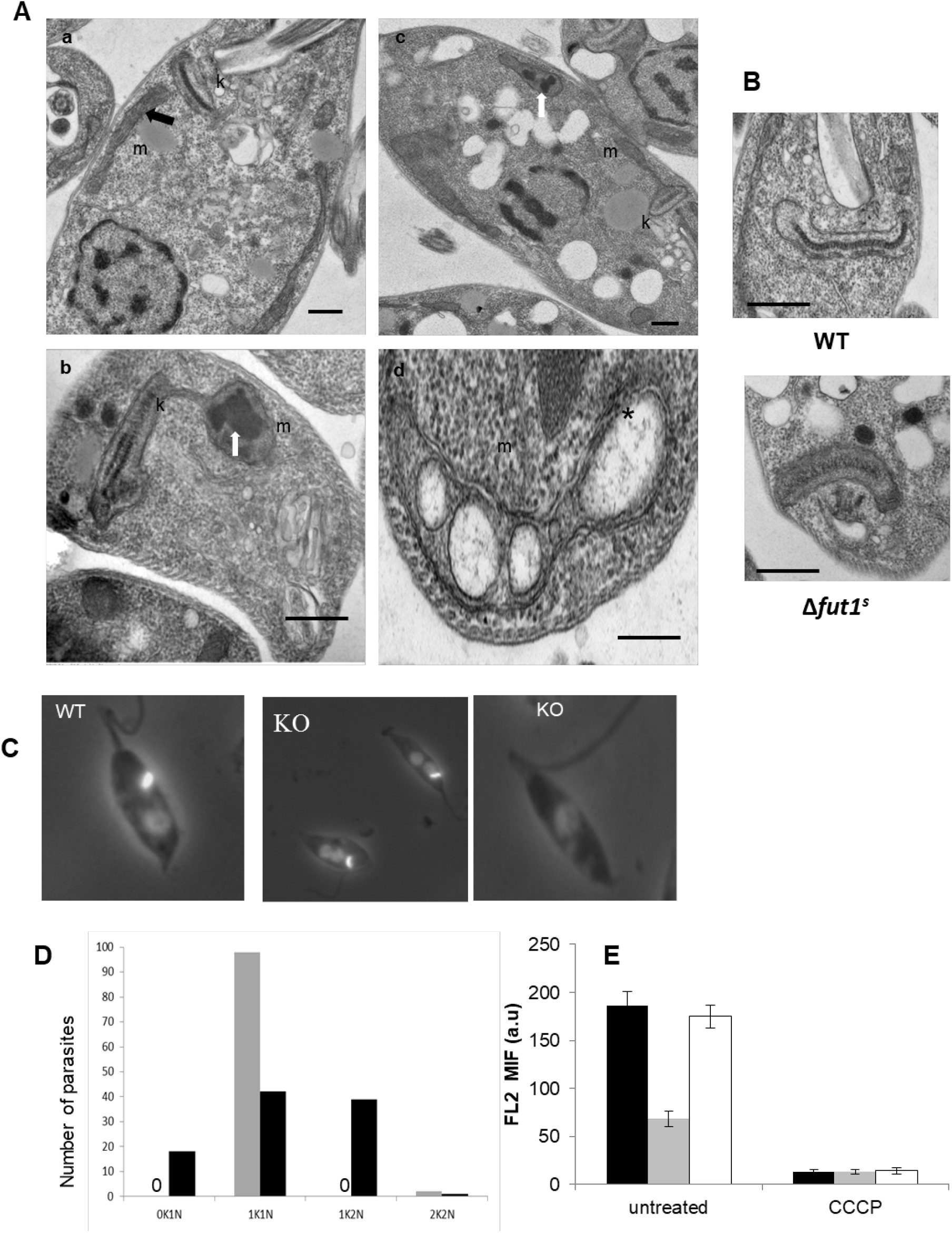
Δ*fut1^s^* shows multiple mitochondrial abnormalities. Panel A, transmission EM of WT (subpanel a) and Δ*fut1****^s^*** (subpanels b-d). m, mitochondria; k, kinetoplast; black arrow, normal mitochondrial cristae; white arrows, aggregates inside mitochondria; star, bloated cristae. Bars = 500 nm. Panel B. Ultrastructural analysis of kDNA in WT and Δ*fut1****^s^***. While WT cell presents typical compact kDNA structure, mutant cell shows a ‘looser’ kDNA network that is wider with decreased length (see Fig. S6 for quantitation). Black bar: 500 nm. Panel C. Loss of kDNA in Δ*fut1****^s^***. WT or Δ*fut1****^s^*** were stained with DAPI to visualize the kDNA network (small bright spot) and nucleus (dimmer large circle). A typical 1 kinetoplast / 1 nucleus (1K/1N) pattern is shown for WT in the leftmost panel; the central panel shows a Δ*fut1****^s^*** cell with a 1K/2N pattern and the rightmost panel shows a Δ*fut1****^s^*** cell lacking kDNA (0K/1N). Panel D. Quantitation of kDNA/nucleus patterns seen in WT (gray bars) vs Δ*fut1****^s^***. (dark bars). A “0” indicates no cells of that pattern Panel E. Mitochondrial potential assessed by staining with TMRE. Parasites were incubated with 100 nM TMRE and analyzed by flow cytometry, with the signal on the FL-2 channel expressed in arbitrary units (a.u.) . The left portion shows cells incubated for 15 min; the right portion shows cells further incubated with 300 µM cyanide m-chlorophenylhydrazone (CCCP) for 60 min. Lines tested are WT, black bar; Δ*fut1****^s^***, gray bar; and Δ*fut1****^s^***/+pXNGPHLEO-FUT1, white bar.

We assessed the mitochondrial membrane potential (ΔΨm) of *Δfut1**^s^*** using tetramethylrhodamine ethyl ester (TMRE), a cationic lipophilic dye whose accumulation depends on mitochondrial membrane potential. Logarithmic phase *Δfut1**^s^*** incubated with TMRE showed only 36% of the WT fluorescence (Fig 8E), which returned to normal upon restoration of *FUT1* (Fig. 8E, panel labelled ‘untreated’). As a control, cells were treated with the protonophore carbonyl cyanide p-trifluoromethoxy-phenylhydrazone (CCCP) to induce loss of mitochondrial membrane potential, which yielded greatly decreased TMRE fluorescence (Fig. 8E, panel labelled CCCP). Thus, loss of *FUT1* results in a profound mitochondrial dysfunction.

### Efforts to detect potential *FUT1*-dependent products in *Leishmania major*

We surveyed unsuccessfully for *FUT1*-dependent products within the parasite mitochondrion, comparing WT and *Δfut1**^s^*** parasites. No mitochondrial specific-labelling was observed using with fluorescent lectins from *Aleuria aurantia* (AAL), *Ulex europaeus* (UEA) or Isolectin B4 (IB4) (57), or with antisera to anti-blood group antigen H type 1 (58) or anti O-Fucβ1-3Glc (L. Zhang and L. Mahal, personal communication). For both lectins and antibodies, often labelling outside of the mitochondrion was observed, possibly arising from one of the other secretory pathway glycosyltransferases (Table 1). Bioorthogonal labelling with 6-alkynyl fucose showed no differences, although labelling was seen in a region in the vicinity of the Golgi/flagellar pocket in both. Such labelling, was absent in the *Δlpg2**^-^*** mutant lacking the Golgi GDP-Man/L-Fuc/D-Arap transporter (59–61), suggesting that 6-alkymyl fucose had been activated to GDP-6-alkynyl-Fuc and trafficked to the secretory pathway for use by FUTs therein (Table 1).

## Discussion

Previous studies have shown that GDP-Fucose synthesis is essential in trypanosomatids, yet for *Leishmania* and *Trypanosoma brucei* fucosylated molecules that might account for this requirement remained unknown. We reasoned that identification of essential fucosyltransferase(s) and characterization of their substrate specificities would better inform our search for the postulated essential fuco-conjugates. Given the structural similarity of L-Fucose and D-arabino*pyranose* and the tendency of enzymes using either substrate (or their activated GDP derivatives) to accept the other, we searched in *Leishmania* genomes for proteins showing relationship to diverse collection of prokaryotic and eukaryotic fucosyl- or D-arabinosyl transferases (Table 1). Four candidates were predicted to be targeted to the secretory pathway, as typical for most parasite glycosyltransferases. *SCA1* and *SCA2* encoded transferases mediating developmentally regulated side-chain D-arabinopyranosylation of LPG within the Golgi apparatus, and unlikely to be essential given that LPG is not (15). The role(s) of *SCAL* and *FUT2* are unknown (Table 1), however, both genes could be readily deleted with little apparent consequence *in vitro* (Figs S1, S2). Amongst possible roles could be D-Ara*p* modification of GIPLs (62), especially as GIPLs are not required for survival (12).

*FUT1* in contrast was conserved in both African and South American trypanosomes, and uniquely, was predicted and then experimentally confirmed to be targeted exclusively to the parasite mitochondrion, most likely within the lumen (Figs. 1, 3). By plasmid segregation tests, *FUT1* appeared to be essential for normal growth (Fig. 2), and as discussed further below, clearly encodes a fucosyltransferase activity. As mitochondrial glycosyltransferases are uncommon, we used a ‘plasmid shuffle’ assay to test whether mitochondrial localization was required by the parasite, examining *FUT1*s with mutated or blocked mitochondrial targeting sequences. Disruption of mitochondrial targeting was found as expected, but these were unable to satisfy the WT *FUT1* requirement (Fig. 6). We similarly explored whether the FUT1 catalytic activity was required for growth, mutating conserved putative active site residues, and again, the mutant enzyme could not satisfy the *FUT1* requirement (Fig. 6). Importantly, all mutant FUT1 proteins were expressed at levels comparable to WT in *vivo* (Fig. 4B), and *in vitro* assays of recombinant enzymes showed similar activities for the WT and mitochondrial targeting mutants, while the catalytic mutant showed near-background activity (Fig. 6E). These data established that both mitochondrial localization and catalytic activity were required for essential *FUT1* function.

One advantage of the plasmid segregation test for essentiality is that it allows tests of many more events than typical transfection based approaches (44). In standard media we tested more than 200 events (Fig. 7), and once FUT1’s mitochondrial role was recognized, we repeated these studies in richer media potentially sparing mitochondrial function. However despite testing more than 1000 estimated events, we were able to obtain only a single mutant completely lacking *FUT1* entirely (*Δfut1**^s^***; Figs. 7, 8). Notably *Δfut1**^s^*** displayed <u>exactly</u> the phenotypes expected for a dysfunctional mitochondrial mutant, including slow growth, structural alterations of cristae, mitochondrial inclusions and abnormalities, alterations and even loss of kinetoplast DNA network, and greatly decreased mitochondrial membrane potential (Figs. 7, 8). Importantly, restoration of FUT1 expression (*Δfut1**^s^/+**FUT1*) completely reversed these defects back to WT. This raises the question as to why *Δfut1**^s^*** was so rare; one explanation is that it bears a second site compensating mutation, albeit one unable to restore normal mitochondrial function. Future studies will address this possibility.

Thus, cell biological, genetic manipulation and the severe mitochondrial dysfunction of a *Δfut1**^s^*** mutant all establish the localization and critical functionality of FUT1 within the parasite mitochondrion. The questions now becomes what is the natural FUT1 acceptor(s) therein, what are the products ultimately requiring FUT1 synthesis, and what role do they play in essential mitochondrial biology? Remarkably, despite knowledge of essential GDP-Fucose synthesis (30–32), and extensive work characterizing parasite glycoconjugates since the early 1980s, few candidate fucosylated molecules have been found in any *Leishmania* species, and none in *L. major*. *L. donovani* expresses a mannose-fucose conjugate whose structure has not been definitively established (33), and several *L. donovani* proteins exhibited MS/MS signatures suggestive of fucosylation (34). Two were mitochondrial HSPs, peptides of which could be fucosylated by FUT1 *in vitro* (Figs. 5B, S5). However in a global proteomic screen identifying nearly 7000 unique proteins, or 80% of the predicted proteome (63), we were unable to identify signs of protein–O-fucose in *L. major* (one caveat is that complex or atypical structures would not have been detected by the MS methodology used). Similarly, comparisons of WT and *Δfut1**^s^*** using a variety of reagents specific for fucoconjugates were uninformative, as was biorthogonal labelling with 6-alkynl fucose. Failure to detect a mitochondrial product might signify a) products below the detection limits, b) inability of modified fucose derivatives such as 6-alkynyl fucose to be properly transported, activated or resist degradation within the mitochondrion (although some data suggested this could occur in the secretory pathway), or c) further incorporation and/or metabolism of fucose to forms not recognizable by the methods used here. A variety of fucose modifications have been described in prokaryotes, and the internal fucose within the SKP1 pentasaccharide D-Galα1-6-D-Galα1-L-Fucα1-2-D-Galα1-3GlcNAc of several single-celled eukaryotes is not recognized by lectins or antibodies tested here (64). Studies searching for the cellular FUT1-dependent products in *T. brucei* have been similarly unrequited (1). Future studies will be necessary to resolve the nature of the GDP-Fucose and *FUT1*-dependent product(s) in trypanosomatids, which genetic and biochemical data strongly predict must nonetheless exist.

We did not deeply explore the glycan acceptor specificity of LmjFUT1, with the best acceptor being LNB and little activity with others tested (Fig. 5B). The presence of significant GDP-Fuc hydrolysis with all substrates tested may be a sign that the native acceptor moiety has yet to be identified (Fig. 5). More extensive studies with the TbFUT1 showed a preference for lac-N-biose and its β-methyl glycoside, again typically accompanied by significant background hydrolysis (1). The strong homology between these two species enzymes and the ability of *TbFUT1* to satisfy the *Leishmania FUT1* requirement (Fig. 5C; (1)) suggests that biologically relevant specificities may be similar in both species. While two substrates for mammalian protein O-fucosyltransferase were inactive with LmjFUT1, two peptides predicted to be fucosylated in *L. donovani* functioned as acceptors, and MS/MS of the one tested confirmed fucosylation (Figs. 5B, S5). Thus LmjFUT1 is unusual in its ability to fucosylate a diverse spectrum of substrates *in vitro*.

While intra-mitochondrial glycosylation is uncommon, that mediated by the mitochondrial isoform of the mammalian O-GlcNAc transferase has received considerable attention (65, 66). Proteomic surveys have revealed approximately 100 O-GlcNAc modified mitochondrial proteins (67–69), mostly encoded by nuclear genes raising the possibility of glycosylation prior to mitochondrial import (70). However, to date no mechanistic studies establish that glycosylated proteins can be imported into mitochondria, while several proteins encoded by mitochondrial DNA are glycosylated (thus rendering cytoplasmic fucosylation unlikely), and mitochondrial transporters able to transport UDP-GlcNAc have been identified (68, 69, 71, 72), in support of the intra-mitochondrial glycosylation paradigm. Importantly, O-GlcNAc modifications of mitochondrial proteins have been postulated to play key roles in mitochondrial energy metabolism and function (68, 69), and an analogous role could be imagined for intra-mitochondrial fucosylation in *Leishmania.* Potentially, trypanosomatid mitochondrial FUT1s may offer a facile system in the future for probing mitochondrial glycosylation in a setting uncomplicated by multiple isoforms targeted to diverse compartments, and its essentiality renders it an attractive target for chemotherapy of trypanosomatid parasites. Indeed, preliminary studies suggest that the *Δfut1**^s^*** mutant cannot survive when inoculated into susceptible murine hosts.

## Materials and Methods

### *Leishmania* culture and transfection

*L. major* strain FV1 (LmjF or WT; WHO code MHOM/IL/80/Friedlin) was grown at 26°C in M199 medium (U.S. Biologicals) containing 10% heat-inactivated fetal bovine serum and other supplements and transfected by electroporation using a high voltage protocol as described (73, 74). Following transfection, cells were allowed to grow for 16-24 h in M199 medium and then plated on semisolid media containing 1% Nobel agar (Fisher) and appropriate selective drugs (30 μg/ml puromycin, 100 μg/ml nourseothricin, 10 μg/ml phleomycin, or 10 μg/ml G418). Individual colonies were picked and grown in liquid medium in same drug concentration as used in plates. Clones were maintained in selective medium for less than 10 passages before use, prior to which selective drug was removed for one passage.

#### Molecular constructions and primers

Molecular constructs are described in detail in Table S1 and oligonucleotide primers are described in Table S2. Molecular constructs were confirmed by restriction mapping, sequencing, and functional testing. Molecular methods were performed as described (30).

### Homologous replacement of chromosomal FUTs, often in the presence of ectopically expressed *FUTs*

(Fig. 2). For *FUT1,* pXNGPHLEO-*FUT1* (B7073) was first transfected to LmFV1 WT, yielding WT / +pXNGPHLEO-*FUT1*. The presence of plasmid was confirmed by PCR and FACS analysis for the presence of *PHLEO* and GFP markers, respectively. The targeting fragments *FUT1*::*SAT* and *FUT1*::*PAC* were liberated from pGEM-*FUT1-SAT* (B7081) or pGEM-*FUT1-PAC* (B7079) by digestion with BamHI and HindIII, and then sequentially transfected into WT/ +pXNGPHLEO-*FUT1* with nourseothricin or puromycin selection, to generate Δ*fut1*::*PAC*/Δ*fut1*::*SAT/ +pXNGPHLEO-FUT1* (termed Δ*fut1***^-^***/ +pXNGPHLEO-FUT1*). For plasmid shuffling experiments, Δ*fut1***^-^***/ +pXNG-FUT1* was further transfected with episomal pIR1NEO (B6483) into which test *FUT1* genes had been inserted (Table S1).

A similar protocol was followed for *FUT2*, substituting pXNGPHLEO-FUT2 and targeting fragments *FUT2*::*HYG* and *FUT2*::*PAC* (Table S1, Fig. S2), to generate Δ*fut2*::*PAC*/ Δ*fut2*::*HYG / +pXNGPHLEO-FUT2*, hereafter termed Δ*fut2* **^-^** */ +pXNGPHLEO-FUT2*. Deletion of both chromosomal copies of *SCAL* was accomplished by successive introduction of targeting fragments *SCAL*::*HYG* and *SCAL*::*BSD* (Table S1; Fig. S1), yielding Δ*scal*::*HYG*/ Δ*scal*::*BSD,* hereafter termed Δ*scal***^-^**.

### Plasmid segregation or shuffling by single cell sorting

Δ*fut1***^-^** parasites bearing pXNGPHLEO-FUT1 alone (segregation) or additionally expressing test FUT1 sequences from pIR1NEO (shuffling) were subjected to flow cytometry and single cell cloning to recover GFP+ cells (bright) or GFP-negative (dim), the latter representing potential segregants or ‘shuffles’. Prior to flow cytometry, cells were incubated in M199 medium without phleomycin for 24 h. washed with phosphate-buffered saline (PBS), and filtered through CellTrics 50 μm filters to remove particulates. Single cell sorting of GFP “bright” or “dim” cells was performed with appropriate gates (Fig. 2C illustrates typical gating settings) using a Dako MoFlo high-speed cell sorter, with single cells selected by stringent gating on forward and side scatter parameters. Single cells were placed into individual wells of 96-well plates, each containing 150 μl M199 medium or Schneider medium. Plates were incubated at 26° C for at least 2 weeks and parasite growth assessed.

### Subcellular localization of FUT1

Logarithmic phase parasites expressing FUT-HA were collected by centrifugation and resuspended in DMEM media containing Mitotracker Red CMXRos (Invitrogen; 50 nM) for 15 min. Parasites were then washed in DMEM and fixed in 4% paraformaldehyde (in PBS) for 5 min, washed in PBS, and deposited on coverslips by centrifugation. The coverslips were then incubated in blocking buffer (PBS containing 0.1% (v/v) Triton-X-100 and 5% (v/v) normal goat sera) for 30 min prior to staining with a rabbit anti-HA polyclonal antibody (Invitrogen, diluted 1:100 in blocking buffer) for 1 h. Following extensive washing in PBS, the coverslips were then stained with an Alexafluor488 goat anti-rabbit antibody (Invitrogen, diluted 1:1000 in blocking buffer).

### Immunoelectron Microscopy

Parasites were fixed in 4% paraformaldehyde/0.05% glutaraldehyde (Polysciences Inc., Warrington, PA) in 100 mM PIPES/0.5 mM MgCl_2_, pH 7.2 for 1 hr at 4°C. Samples were then embedded in 10% gelatin and infiltrated overnight with 2.3 M sucrose/20% polyvinyl pyrrolidone in PIPES/MgCl_2_ at 4°C. Samples were trimmed, frozen in liquid nitrogen, and sectioned with a Leica Ultracut UCT cryo-ultramicrotome (Leica Microsystems Inc., Bannockburn, IL). 50 nm sections were blocked with 5% FBS/5% NGS and subsequently incubated with anti-HA antibody followed by streptavidin conjugated to 15 nm colloidal gold (BB International, Cardiff, UK). Sections were washed in PIPES buffer followed by a water rinse, and stained with 0.3% uranyl acetate/2% methyl cellulose. Samples were viewed with a JEOL 1200EX transmission electron microscope (JEOL USA Inc., Peabody, MA). Parallel controls omitting the biotinylated agglutinin were consistently negative at the concentration of streptavidin used in these studies. For quantitation, images from sections intersecting the mitochondrion, cytosol and nucleus were chosen; for each beads were counted in each compartment, whose area was measured using Image J. The bead counts were normalized to the relative area of each compartment measured across all sections (1:3:2 for mitochondrion: cytosol: nucleus).

### Transmission electron microscopy

Parasites were fixed with 4% paraformaldehyde (PFA) and 2% glutaraldehyde in 0.1 M phosphate buffer (pH 7.4) for 4 h at room temperature (RT). The samples were then washed in 0.1 M phosphate buffer, postfixed in 2% OsO4, and encapsulated in agarose. This was followed by dehydration in ascending grades of ethanol, infiltration, embedding in an Epon 812-araldite plastic mixture, and polymerization at 60°C for 24 h. Ultrathin sections (50 to 70 nm) were obtained using an ultramicrotome (Leica Ultracut UCT; Leica Microsystems GmbH, Wetzlar, Germany) and picked up onto 200 mesh copper grids. The sections were double stained with uranyl acetate and lead citrate and observed under a FEI Tecnai-12 twin transmission electron microscope equipped with a SIS MegaView II CCD camera at 80 kV (FEI Company, Hillsboro, OR, USA).

### Measurement of mitochondrial membrane potential (ΔΨm) in live Leishmania promastigotes

Tetramethylrhodamine ethylester perchlorate (TMRE, Molecular Probes, Inc., Eugene OR, USA) was prepared at a concentration of 1 mM in DMSO and stored at -20°C. *L. major* promastigotes were washed once in PBS, resuspended at 2×10^6^ cells/ml in PBS containing 100 nM TMRE, incubated for 15 min at room temperature, and analyzed by flow cytometry on the FL2-H channel. Baseline TMRE was recorded and expressed as mean fluorescence in arbitrary units (a.u.). Subsequently, carbonyl cyanide m-chlorophenylhydrazone (CCCP, Sigma) was added to the cells (300 mM final concentration) and TMRE fluorescence was measured at various times up to 60 min.

### Protein expression and purification

GST- and His-tagged recombinant *FUT1* constructs were transformed to BL21 (DE3) *E. coli* strain. Cells were grown at 37 °C until OD_600_ reached about 0.5, when 50 µM isopropyl-β-D-thiogalactopyranoside (IPTG) and incubated at 15 °C for 16 h. Cells were harvested by centrifugation and incubated 1 hr on ice in 10 mM TrisHCl, pH 7.5, 0.15 M NaCl, 1 mM DTT, 1 mg/ml lysozyme (Sigma) with EDTA-free Complete Protease Inhibitor Tablet (Roche), before disrupting by sonication. The soluble fraction was obtained by centrifugation at 17,000 × *g*, 4 °C for 30 min. For GST-tagged protein purification, the supernatant was incubated with Glutathione Sepharose 4B resin (GE Healthcare) for 1 h at 4 °C. The bound protein was washed with 10 bed columns of 10 mM TrisHCl, 0.15 M NaCl followed by elution using 50 mM Tris, 50 mM reduced glutathione, 0.15 M NaCl, 5 mM DTT, 0.1% Triton X-100, pH 8.5. For His-tagged PTR1-FUT1, supernatant was incubated with Ni-NTA resin (Qiagen) for 1 h at 4 °C. The bound protein was washed with 10 bed columns of 10 mM TrisHCl, pH 8.0, 0.15 M NaCl, 50 mM imidazole followed by elution using 10 mM TrisHCl, M NaCl, 250 mM imidazole, 5 mM DTT, 1% Triton X-100. The yield of the recombinant protein was estimated by Qubit 2.0 (Invitrogen) and the degree of purification by SDS-PAGE analysis.

### Fucosyltransferase activity assays

Enzymatic activity was determined by GDP-Glo bioluminescent GDP detection assay with GDP-Fucose (Promega Corporation, Madison, WI USA). Aliquots of 3 µg of the affinity purified recombinant proteins were incubated with 600 µM GDP-Fuc, 10 mM acceptor (β-D-Gal-(1→3)-D-GlcNAc; lac-N-biose or LNB), N-acetyllactosamine (LacNAc), β-Galactose, or Galβ1,3GalNAc) or 1.7 mM peptide from mHSP70 or mHSP60 (EWKYVSDAEKE or SKELESLANDS, respectively; Peptide2go, Manassas, VA) in 20 mM sodium phosphate, 25 mM KCl, 5 mM MgCl_2,_ pH 6.0 or 40 µM acceptor (TSR or EGF repeats, provided by R. Haltiwanger, U. Ga.) in 50 mM imidazole-HCl (pH 7.0), 50 mM MnCl_2_, in a final volume of 25 µl for 5 min at 37 °C. Reactions were terminated by boiling for 5 min before 25 µl of GDP detection regent was added and incubated at RT for 1 h. Luminescence was recorded using a Microplate Luminometer or IVIS imaging system (Perkin Elmer, IL)

For peptide mass spectrometry, a reaction mix of LmFUT1 with the mHSP70 peptide above was acidified with trifluoroacetic acid (TFA) to a final concentration of 1%. Peptides were bound to porous graphite carbon micro-tips (Glygen) (PMID 22338125) and were eluted with 60% acetonitrile in 0.1% TFA, and dried in a Speed-Vac after adding TFA to 5%. and dissolved in 2.7 µL of 1% formic acid. Samples were analyzed by LC-MS, using an 75 µm i.d. × 50 cm Acclaim^®^ PepMap 100 C18 RSLC column (Thermo-Fisher Scientific) equilibrated with 1% formic acid and a gradient from 1% formic acid to acetonitrile/1% formic acid and a Q Exactiv™ Quadrupole-Orbitrap™ mass spectrometer. The acquisition of the HCD spectra were triggered from the values corresponding to the expected charge states of the doubly and triply charged protonated molecular ions for the unmodified and fucosylated EWKYVSDAEKE peptide (*m/z* = 692.325, 461.888, 765.354 and 510.572). The MS1 scan was acquired in the Orbitrap mass analyzer with an AGC target value of 3e6 over a scan range of *m/z* = 150 – 2000 at a mass resolving power set to 70,000 (at *m/*z = 200). The unprocessed LC-MS data were analyzed using SKYLINE (version 3.6.9).

### Enzymatic assay of HA-tagged FUT1 expressed in *Leishmania* and bound to anti-HA beads

Assays and resolution of products by HPTLC were performed as described (1). Briefly, *Leishmania* lysates were incubated with Anti-HA-beads in the presence of in the presence of 1 μCi GDP[^3^H]Fuc (American Radiochemicals) and 1 mM LNB in 50 mM TrisHCl, 25 mM KCl, pH 7.2 in a final volume of 25 μl for 2 h at 37°C. Reactions were then cooled on ice, water added to 200 μl final volume, and desalted on a mixed-bed column made of 100 μl each Chelex100 (Na^+^) over Dowex AG50 (H^+^) over Dowex AG3 (OH^-^) over QAE-Sepharose A25 (OH^-^). The products were freeze dried, resuspended in water and dried in a Speedvac concentrator, and dissolved in 20% 1-propanol (HiPerSolv Chromanorm, VWR). They were then separated by HPTLC using 10 cm Si-60 plates (Merck) and separated using 1-propanol : acetone : water 9:6:4 (v:v:v) as mobile phase. Plates were sprayed with En^3^hance® (PerkinElmer) and the materials visualized by fluorography at -80°C using Biomax XAR film (Kodak) with an intensifying screen.

### Lectins, antibodies, and 6-alkynyl labelling

For lectin-mediated imaging, WT, *Δfut1****^s^,*** or *Δlpg2^-^* parasites were grown in the presence of 100 µM to 1mM L-fucose. After 3 days, 2 x 10^6^ parasites were washed, fixed onto glass coverslips, and incubated with 17 µg/mL fluorescein labeled *Aleuria aurantia* lectin (AAL; Vector Laboratories FL-1319), 20 µg/mL *Ulex Europaeus* Agglutinin I (UEAI; Vector Laboratories FL-1061), or 5µg/mL *Griffonia simplicifolia* IB4-Alexa 488 lectin (Thermo Fisher; kindly provided by Igor Almeida) at room temperature in the dark for one hour. For antibody-mediated imaging, fixed parasites were incubated with 5µg/mL Blood Group Antigen H (O) Type 1 monoclonal antibody (Invitrogen 13-9810-82), washed three times with 1x PBS and incubated with 1:100 α-mouse secondary antibody conjugated to Alexa 488 (Invitrogen, A11029) each for 1 hour at room temperature. All imaging was performed using a Zeiss LSM880 Confocal Laser Scanning Microscope with Airyscan. For biorthogonal labeling, parasites were inoculated into M199 medium at 1 x 10^5^/mL (WT or *Δlpg2^-^*) or 3.8 x 10^6/^ml (*Δfut1****^s^***) and grown for three days in the presence/absense of 100µM alkyne fucose (Invitrogen C10264), after which parasites were washed in PBS. Prior to confocal microscopy, click labeling was done using azide-488 (Invitrogen A10266) and the Click-iT™ Cell Reaction Buffer Kit (Invitrogen C10269). For western blotting, 1 x 10^6^ labelled parasites were lysed 10 mM Tris pH7.5, 150 mM NaCl, 1% NP-40, and Roche cOmplete EDTA free proteinase inhibitor cocktail (11836170001). Click labeling with biotin azide (Invitrogen B10184) was performed using the Click-iT™ Protein Reaction Buffer Kit (Invitrogen C10276). Proteins were separated using a 10% SDS-PAGE gel and transferred onto a PVDF membrane which was subsequently probed with 1:5,000 fluorescent streptavidin probe (Li-Cor 925-32230). Western blotting with anti-GlcFuc antibody (kindly provided by Lara Mahal) was performed with lysed samples from WT, *Δfut1****^s^,*** or *Δlpg2^-^* parasites grown with or without 1mM fucose as described above; a positive control of 336 ng BSA-GlcFuc was included. The membrane was incubated in using 1µg/mL anti-GlcFuc for 1 hour at room temperature, washed, and probed with 1:5,000 donkey anti-chicken secondary antibody for 1 hour at room temperature (Li-Cor 925-68075). All western blot imaging was performed using the Li-Cor system.

## Acknowledgements

We thank Robert Haltiwanger for providing EGFR and TSR substrates, L. Mahal and L. Zhang for anti-Glc-Fuc antibody, Giulia Bandini for discussions and encouragement, Deborah E. Dobson and Lon-Fy Lye for comments on the manuscript, Wandy Beatty of the Molecular Microbiology Imaging Facility for transmission and immuno-electron microscopy, I.C. Almeida and S. Portillo for providing IB4 lectin, and R.R. Townsend, P.E. Gilmore, and R.W. Sprung of the Washington University Proteomics Shared Resource for performing mass spectrometric analysis. For single cell sorting we thank the staff of the high-speed sorter core, Alvin J. Siteman Cancer Center, Washington University Medical School. This work was supported by NIH grants AI31078 and AI29646 to SMB, an administrative supplement to AI31089 to GP, Berg Postdoctoral Fellowships from the Dept. of Molecular Microbiology (SM, HG), and a Wellcome Investigator Award (101842/Z/13/Z) to MAJF.

---

**Figure S1.**
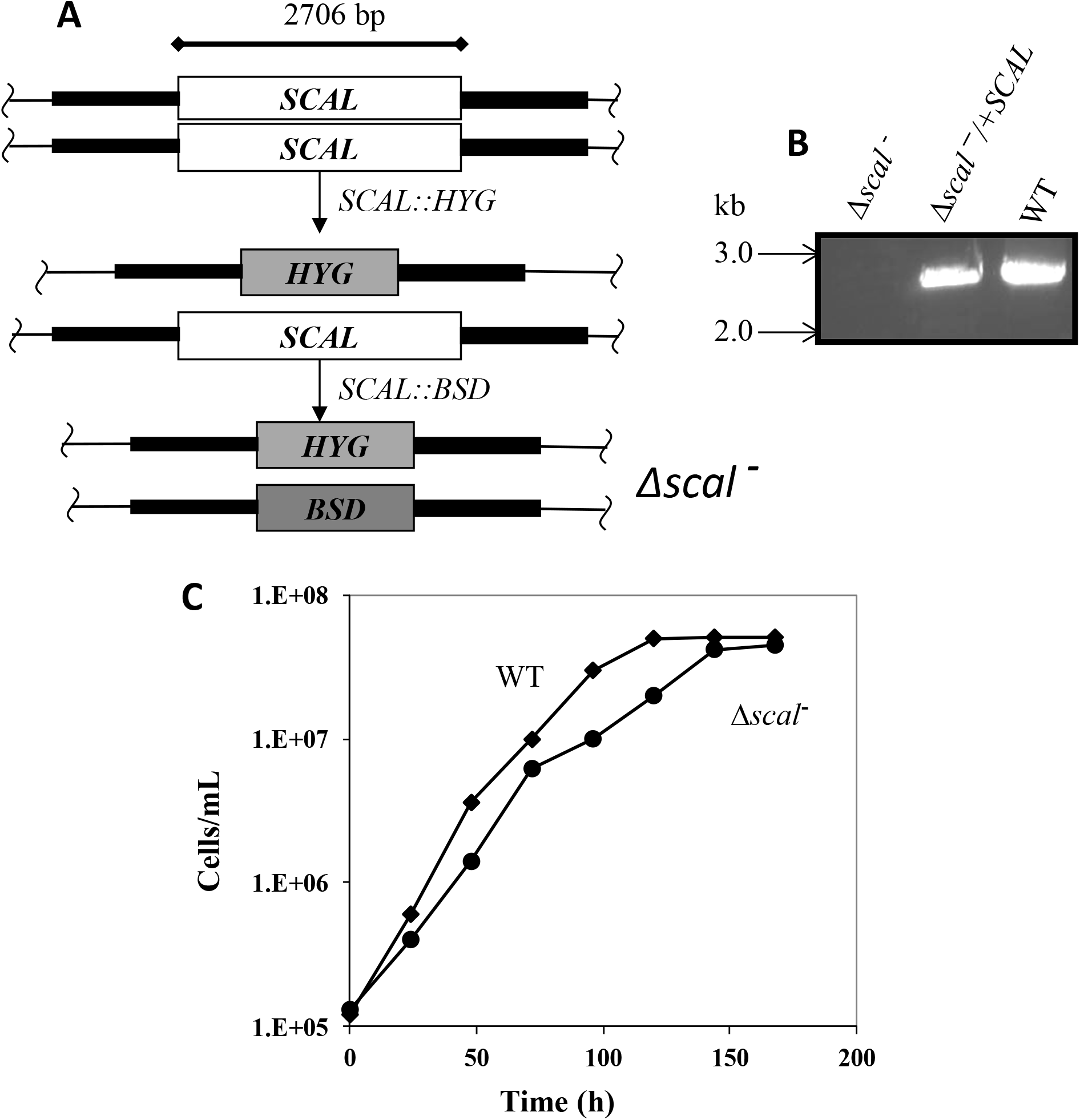
The *SCAL* gene is not essential in *L. major* promastigotes. A. Schematic of the *SCAL* locus showing flanking regions used for homologous gene replacement (dark flanking bars), the SCAL ORF used for PCR tests with primers 3840/3841, and the *SCAL::HYG* or *SCAL::BSD* drug resistance cassettes. WT *L. major* were successively transfected and selected for the *SCAL*::*HYG* and *SCAL*:*BSD* targeting fragments, leading to complete loss of *SCAL (Δscal****^-^***). B. PCR with *SCAL* ORF primers showing loss of *SCAL* in *Δscal* ***^−^*** and retention in WT. and restoration in *Δscal* ***^-^****/ +*pXNGSAT-SCAL C. Growth of WT and Δ*scal*^-^ in M199 medium.

**Figure S2.**
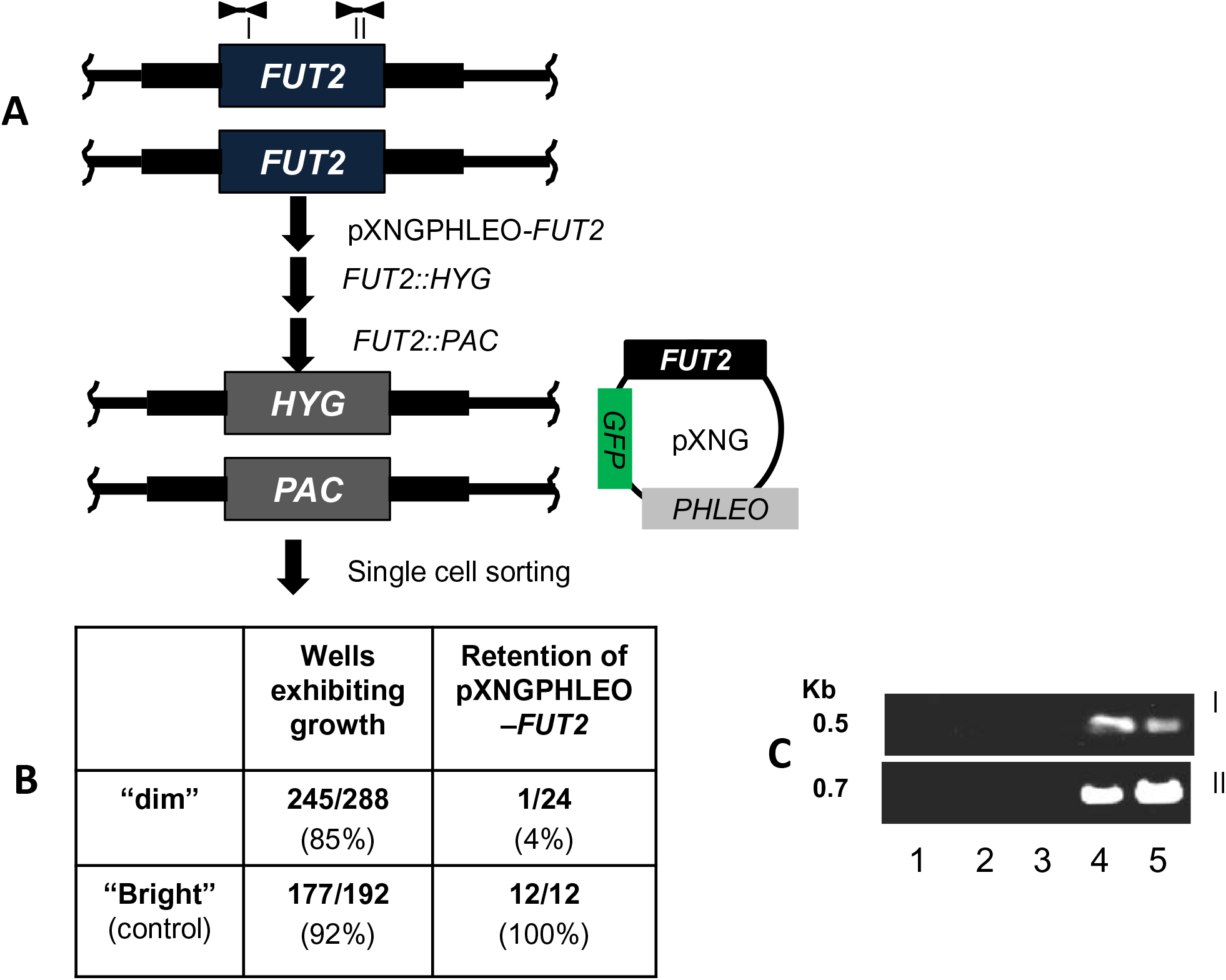
The *FUT2* gene is not essential in *L. major* promastigotes. A. Diagram of the *FUT2* locus and PCR primers used, and the scheme for introduction of episomal FUT2 followed deletion of both chromosomal alleles. Heavy black bars depict the flanking sequences used in targeting constructs. B. Results of plasmid segregation tests; survival of “dim” cells and loss of pXNGPHLEO-*FUT2* established the viability of the Δ*fut2***^-^** cells. C. PCR to confirm loss of all *FUT2* sequences in Δ*fut2***^-^** . Lanes 1-3, 3 different Δ*fut2***^-^** clones; lane 4, Δ*fut2***^-^** /+pIR1SAT-FUT2; lane 5, WT. Primer pairs I (3990/3839) and II (3957/3958) probe the left and right ends of the *FUT2* ORF.

**Fig. S3.**
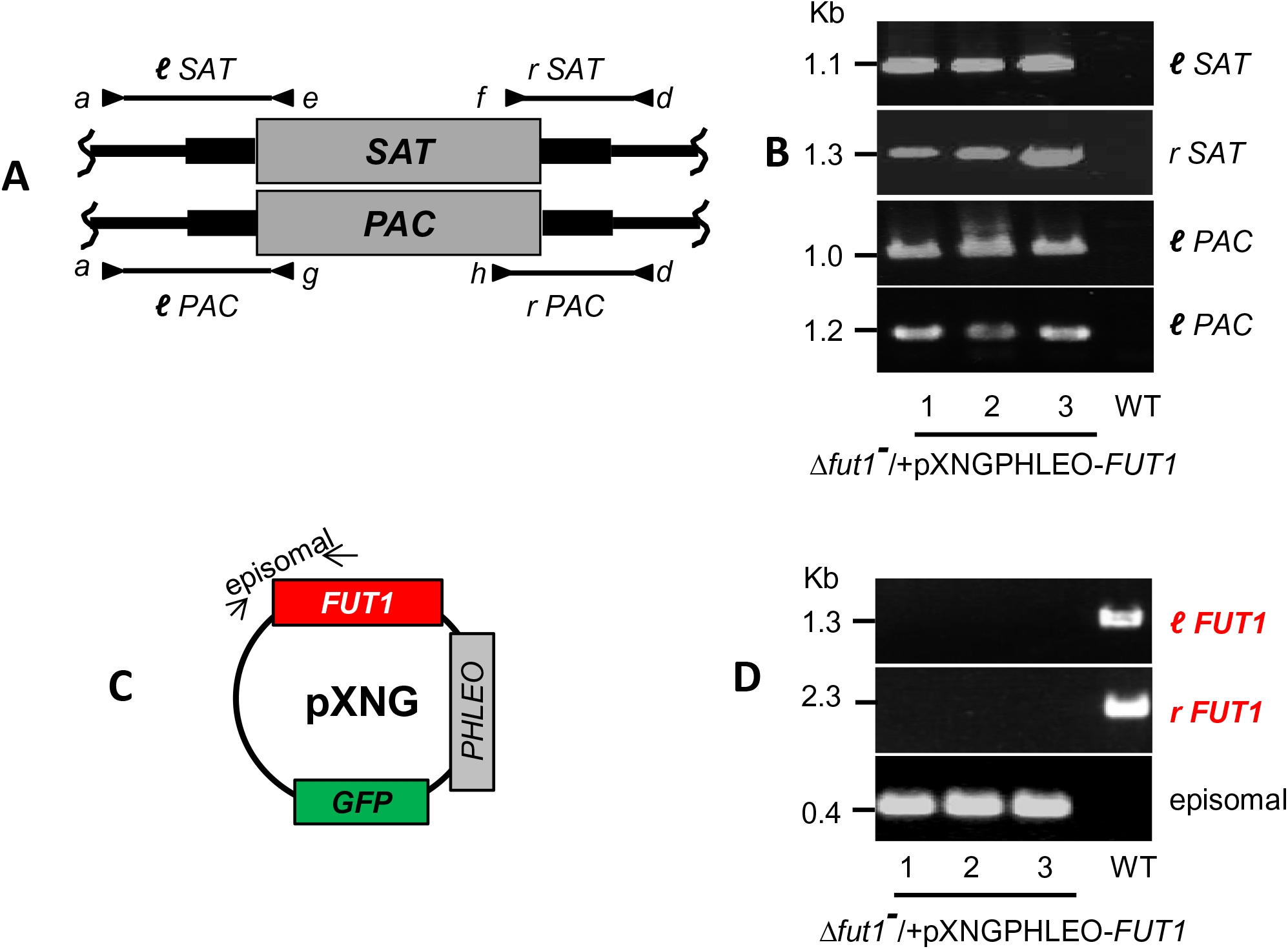
PCR verification tests of *fut1^-^*/+pXNGPHLEO-*FUT1*. A. Schematic of chromosomal loci with *FUT1:SAT* and F*UT1:PAC* replacements. The location of PCR primer pairs used to establish both planned chromosomal replacements is shown. Black bars represent flanking regions used for homologous recombination. B. PCR tests of planned SAT and PAC replacements depicted in panel A. Lanes 1-3 contain three different Δ*fut1^-^*/+pXNGPHLEO-FUT1 clones; lane 4, WT. C. Schematic of the pXNGPHLEO-FUT1 episome. The location of the episomal-specific FUT1 PCR primers is indicated. D. PCR tests confirming expected presence or absence of pXNGPHLEO-FUT1; lanes and samples are the same as in panel B.

**Figure S4.**
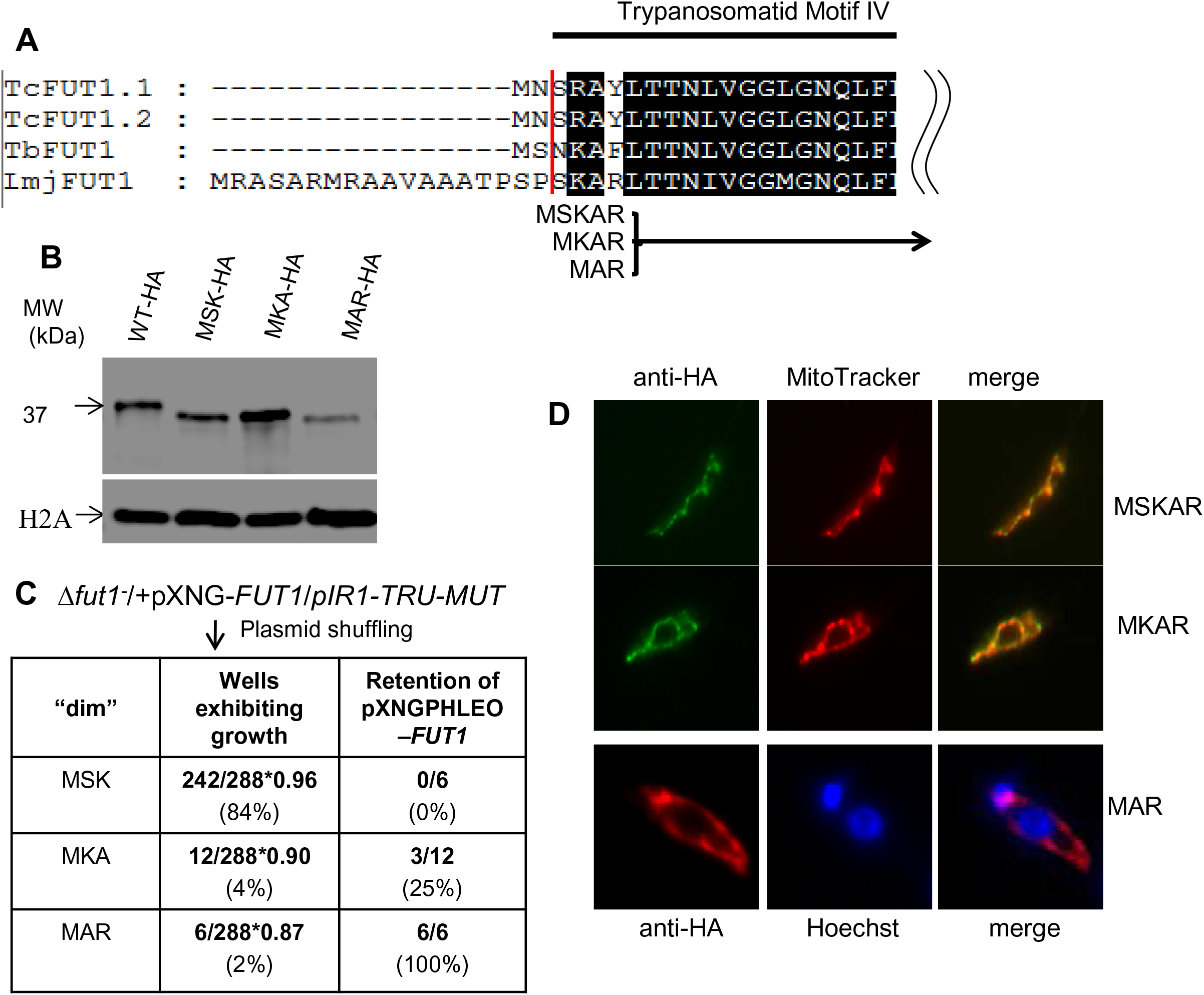
Functionality and mitochondrial localization of N-terminal deletions of LmFUT1. Panel A. Alignment of N-termini of FUTs from Trypanosomatids and scheme of truncated N-terminal LmjFUT mutants (TRU). Panel B. Western blot analysis with anti-HA antisera of whole cell lysates from tagged truncated FUT1 mutants (Δ*fut1^-^* /+pXNG-FUT1/pIR1-TRU-HA). Lane 1: WT-HA, Lane 2: MSK-HA, Lane 3: MKA-HA, and lane 4: MAR-HA. Panel C. Plasmid shuffling to test the function of truncated LmjFUT1s (TRU). Panel D. Indirect immunofluorescence of truncated LmjF mutants described in panel C incubated with anti-HA antibody (green or red, column a, Mitotracker Red CMXRos (red, column b for MSKAR and MKAR mutants), or Hoechst dye (blue, column b for MAR mutant); merged fluorescence images are shown in panel c. For MSKAR and MKAR mutants the pearson’s r colocalization value with the mitochondrion was 0.95 and 0.93.

**Figure S5.**
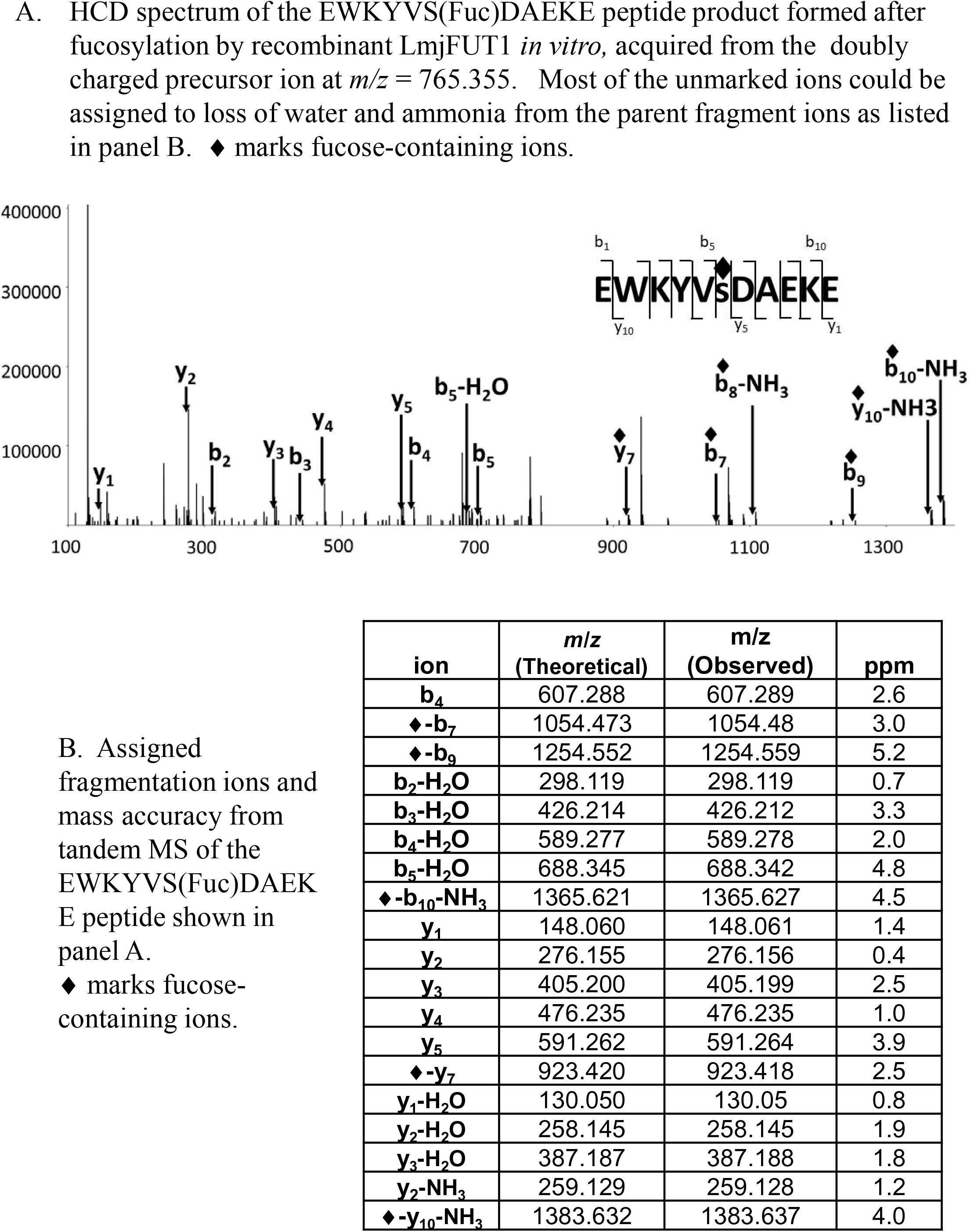
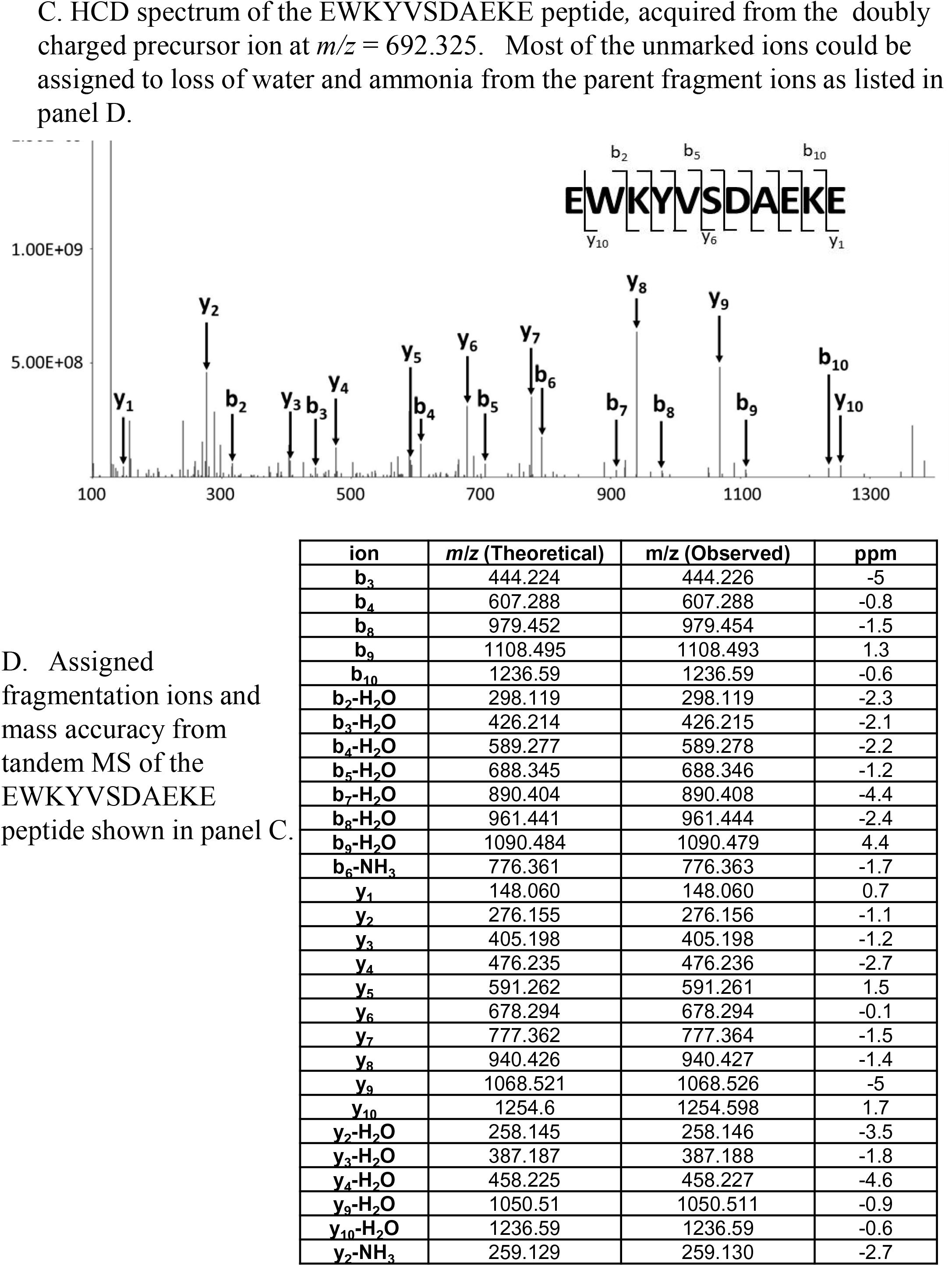
Tandem MS analysis

**Figure S6.**
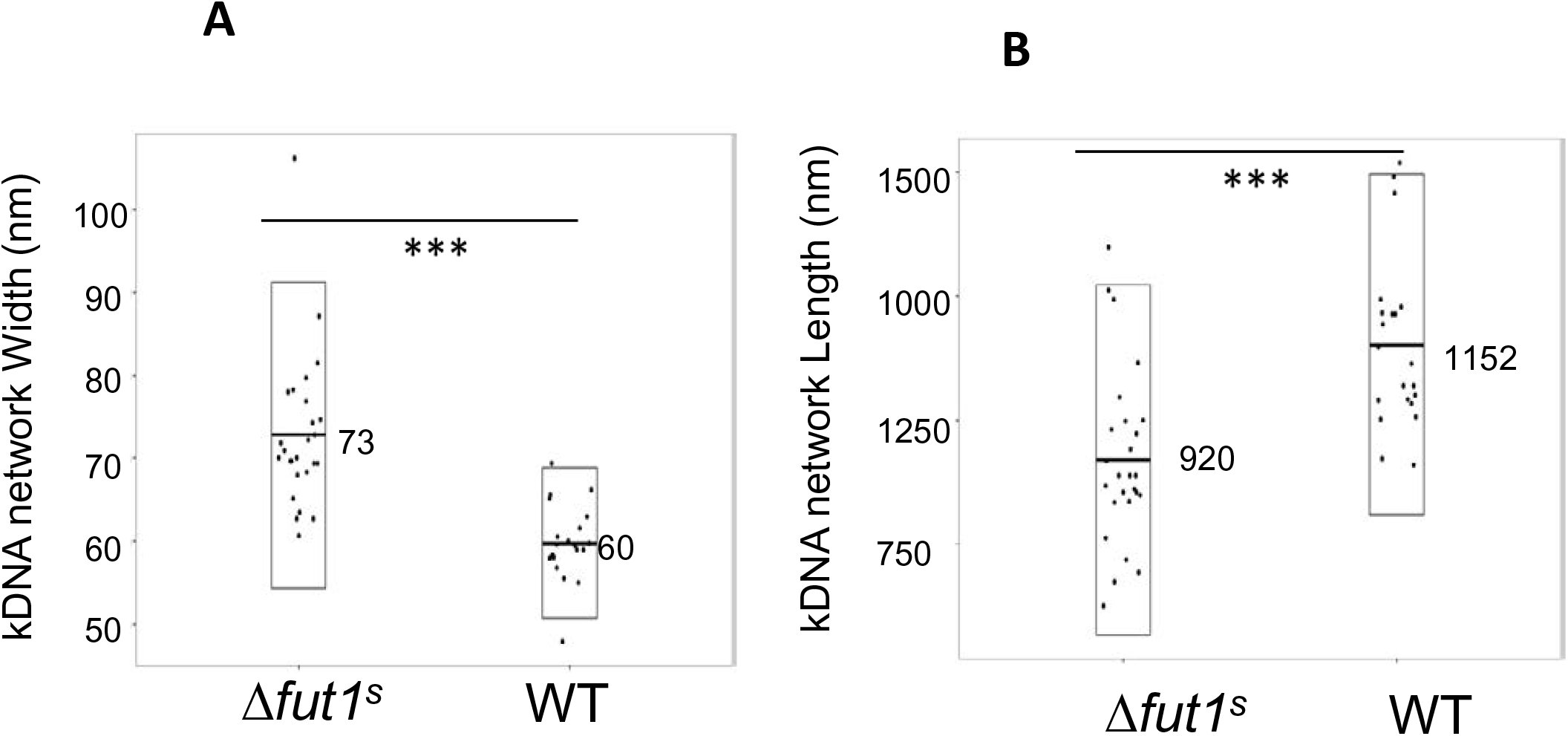
Measurement of the kDNA network dimensions in WT and Δ*fut1^s^.* The kDNA network width and length were measured from transmission EM images using Image J; each dot represents one network. A, width; B, length. ***, *p* value < 0.001

**Table S1.**
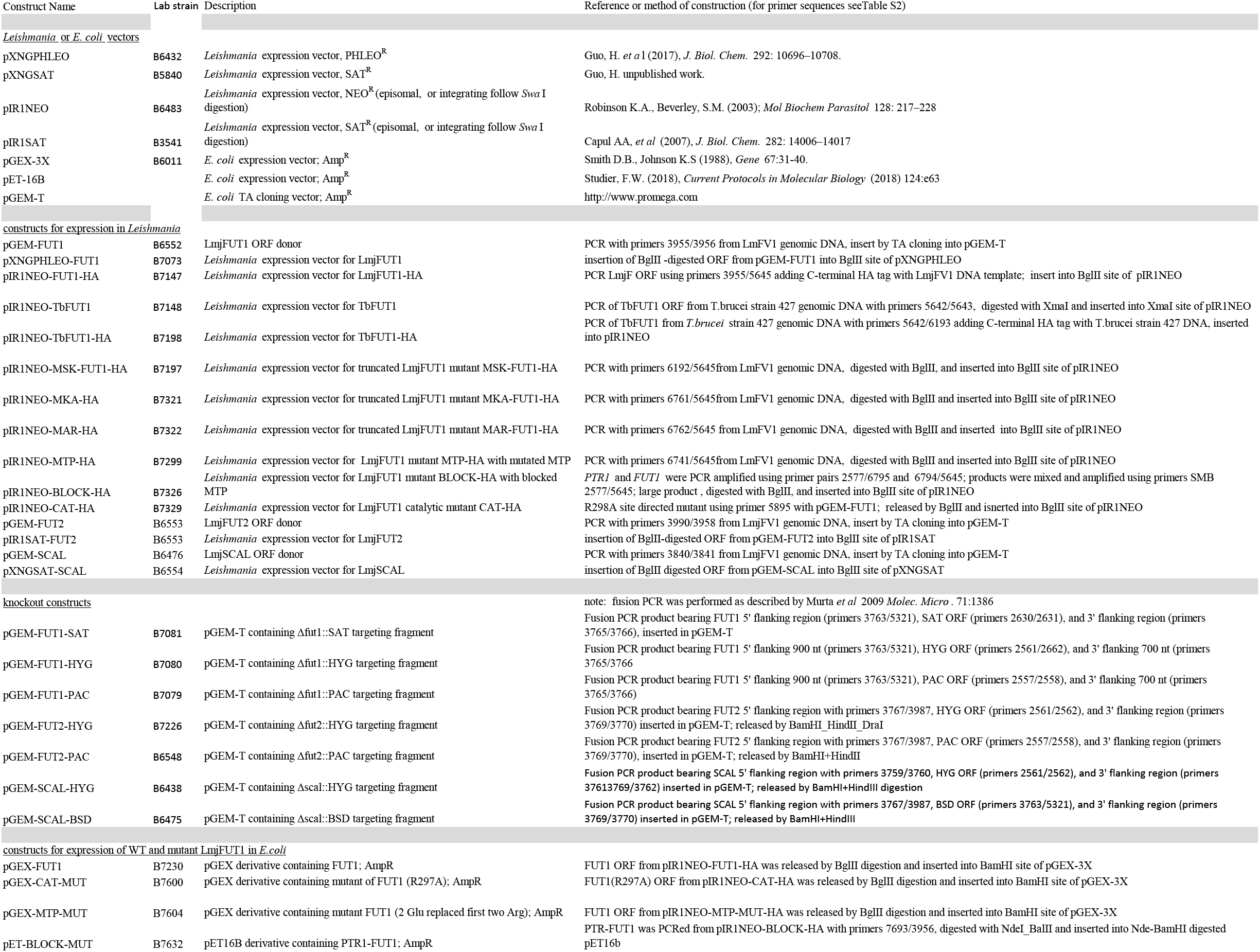
Expression and gene targeting constructs used in this study

**Table S2.**
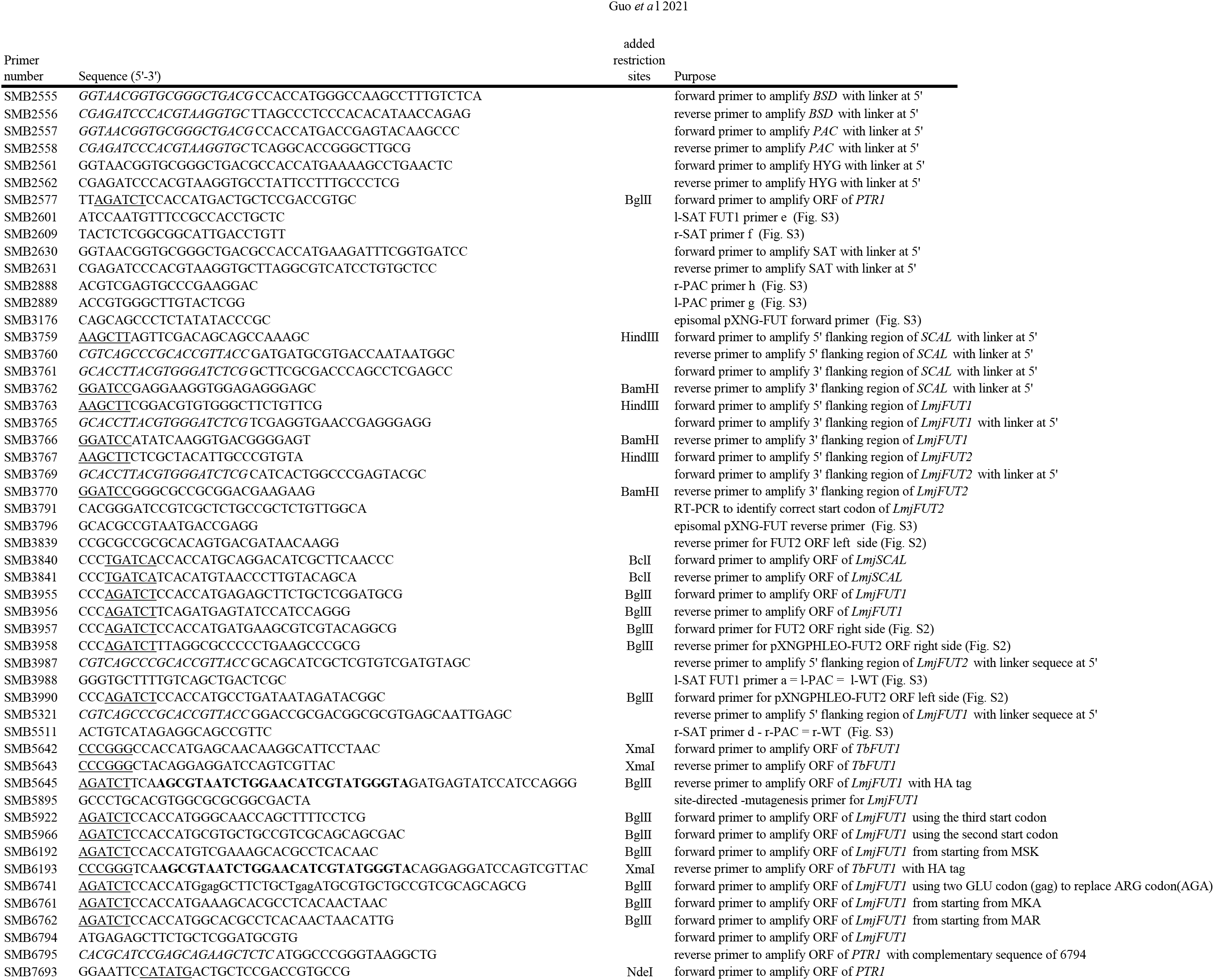
Oligonucleotide primers

